# Integrating Hydrogen Exchange with Molecular Dynamics for Improved Ligand Binding Predictions

**DOI:** 10.1101/2025.01.13.632795

**Authors:** Benjamin T. Walters, Alexander W. Patapoff, James A. Kiefer, Ping Wu, Weiru Wang

## Abstract

We introduce Hydrogen-Exchange Experimental Structure Prediction (HX-ESP), a method that integrates hydrogen exchange (HX) data with molecular dynamics (MD) simulations to accurately predict ligand binding modes, even for targets requiring significant conformational changes. Benchmarking HX-ESP by fitting two ligands to PAK1 and four ligands to MAP4K1 (HPK1), and comparing the results to X-ray crystallography structures, demonstrated that HX-ESP successfully identified binding modes across a range of affinities significantly outperforming flexible docking for ligands necessitating large conformational adjustments. By objectively guiding simulations with experimental HX data, HX-ESP overcomes the long timescales required for binding predictions using traditional MD. This advancement promises to enhance the accuracy of computational modeling in drug discovery, potentially accelerating the development of effective therapeutics.

## Introduction

Structural insights into the interactions between a small molecule and a target protein accelerate modern drug discovery efforts primarily by guiding the affinity maturation process through computational and rational structure-based drug design (SBDD)^1,2^. Traditionally tools like X-ray crystallography and cryo-EM provide these data, but enabling those methods can be resource intensive, particularly for dynamic targets. Pharmacophore modeling and virtual screening^3^, for example, can identify novel scaffolds that can be further developed into high affinity drug candidates in the absence of structure; however, both benefit greatly from information about the binding site. As a result, the need for reliable and accurate computational prediction of ligand binding models remains paramount.

Molecular docking^4–8^ has shown promise in the absence of bound structural data but success typically requires a well-formed binding site in the protein prior to docking^9^. These methods selectively limit the flexibility of the protein (termed the “receptor”) and often the small molecule (called the “ligand”) for computational efficiency making accurate predictions of the binding pose impossible when large changes in the receptor are required^10,11^. The most advanced induced-fit docking programs, like GLIDE, HADDOCK, and AutoDock Vina, address this problem by allowing limited sidechain and backbone flexibility^4^ but selecting which atoms to restrain and which ones to allow motion is highly subjective^12^. Ensemble based docking methods^13^ partially circumvent the issue by docking into multiple receptor conformations; however, generating the appropriate ensemble remains a formidable challenge and is critical for success. Furthermore, the choice of scoring function to evaluate candidate poses introduces another layer of subjectivity. Consensus polling of multiple docking programs^4^ offers advantages in this regard but the problem remains - all docking algorithms ultimately prioritize computational efficiency often at the expense of accuracy.

Alternatively, long molecular dynamics (MD) simulations address the problem by brute-force^14–16^, frequently employing the ligand at high concentration, as a cosolvent, and accurately predicting the bound state where larger conformational changes are required; however, computational time and cost escalate as affinity decreases. Consequently, this approach becomes less tractable for early discovery stages where lower affinity ligands are common. In the later stages of discovery where affinities are improved, experimental structures are often readily available, reducing the need for computational models of ligand binding.

This reality motivated the development of a method that melds experimental hydrogen exchange (HX) data with iterative molecular dynamics coined: Hydrogen-Exchange Experimental Structure Prediction (HX-ESP). HX experiments measure the exchange rate of backbone amide protons and provide valuable structural information about protein conformation and dynamics, making them a promising tool for elucidating where a ligand binds. By integrating HX measurements with iterative molecular dynamics, HX-ESP aims to accurately predict bound ensembles, even when significant conformational changes in the apo receptor are necessary for ligand binding. The HX-ESP algorithm objectively determines how to guide simulations from experimental measurements, thereby eliminating bias and increasing robustness in its use for computational modeling.

We benchmark its performance by fitting two ligands to PAK1 and four ligands to MAP4K1 (HPK1) and comparing results to X-ray structures that were solved in parallel. Known PAK1 ligands bind with minimal protein motion, making docking straightforward; whereas, large conformational changes must occur in HPK1, presenting a significant challenge. The study compares the performance of HX-ESP against HADDOCK (High Ambiguity Driven protein-protein DOCKing), an industry standard for data-guided, induced-fit docking. The results demonstrate that HX-ESP robustly identifies binding modes for ligands, from nanomolar to micromolar affinity, even when conformational changes far beyond the reach of flexible docking are required. This advancement holds promise for improving the accuracy of ligand binding predictions in drug discovery, potentially accelerating the development of effective therapeutics.

### HX-ESP Method Overview

#### 1. Protection Factor Maps and Theory

Amide HX experiments measure the exchange rate of backbone amide protons and have become a popular method to provide structural information about the solution state of proteins^17–19^. Quantifying the protection factor (PFs), or the ratio of observed exchange rates between two states, *PF* = *k_ex_*/*k*′_*ex*_, from HX experiments provides unparalleled access to sub-global thermodynamic and kinetic information at the resolution of individual amino acids^20^. Initially, these measurements were generally applied to questions involving protein folding and local stability ^21,22^ (e.g. where *k*_*ex*_ and *k*′_*ex*_ represent exchange rates from folded and unfolded states). HX experiments are also increasingly used to map where a compound binds to a protein target, capturing both direct and allosteric, or long-range effects that accompany binding^23,24^. In this application, PFs give the work done on the protein by ligand binding: Δ*G*= −*RTln*(*k*_*ex*_/*k*′_*ex*_); *w*= −Δ*G* where *k*_*ex*_ and *k*′_*ex*_ refer to the unliganded and liganded exchange rates, respectively, and *w*represents work. Noting by these relationships that the PF monitors work done to the protein through binding, we can show by the physical principle of efficiency and loss that the binding site must be in proximity to the largest PF(s) measured upon compound addition. This principle holds that useful energy, capable of doing work, must be lost with each successive energy transfer from the source. In the context of ligand binding, each successive transfer potentially occurs with each additional shell of atoms as we move from the Ligand:protein intermolecular contacts. Though small PFs may be in close proximity to the ligand due to variable transfer efficiencies, this principle of loss means that the largest work, or PF, can only be produced from direct interaction with the ligand, and must decrease with distance from the binding pocket. Based on this idea, the first step of HX-ESP involves transforming a PF map (**Figure 1A**) into a perturbation set (PBS, **Figure 1B**), or list of residues most likely to be in close proximity of the ligand on the basis of relative PF magnitude.

**Figure 1.**
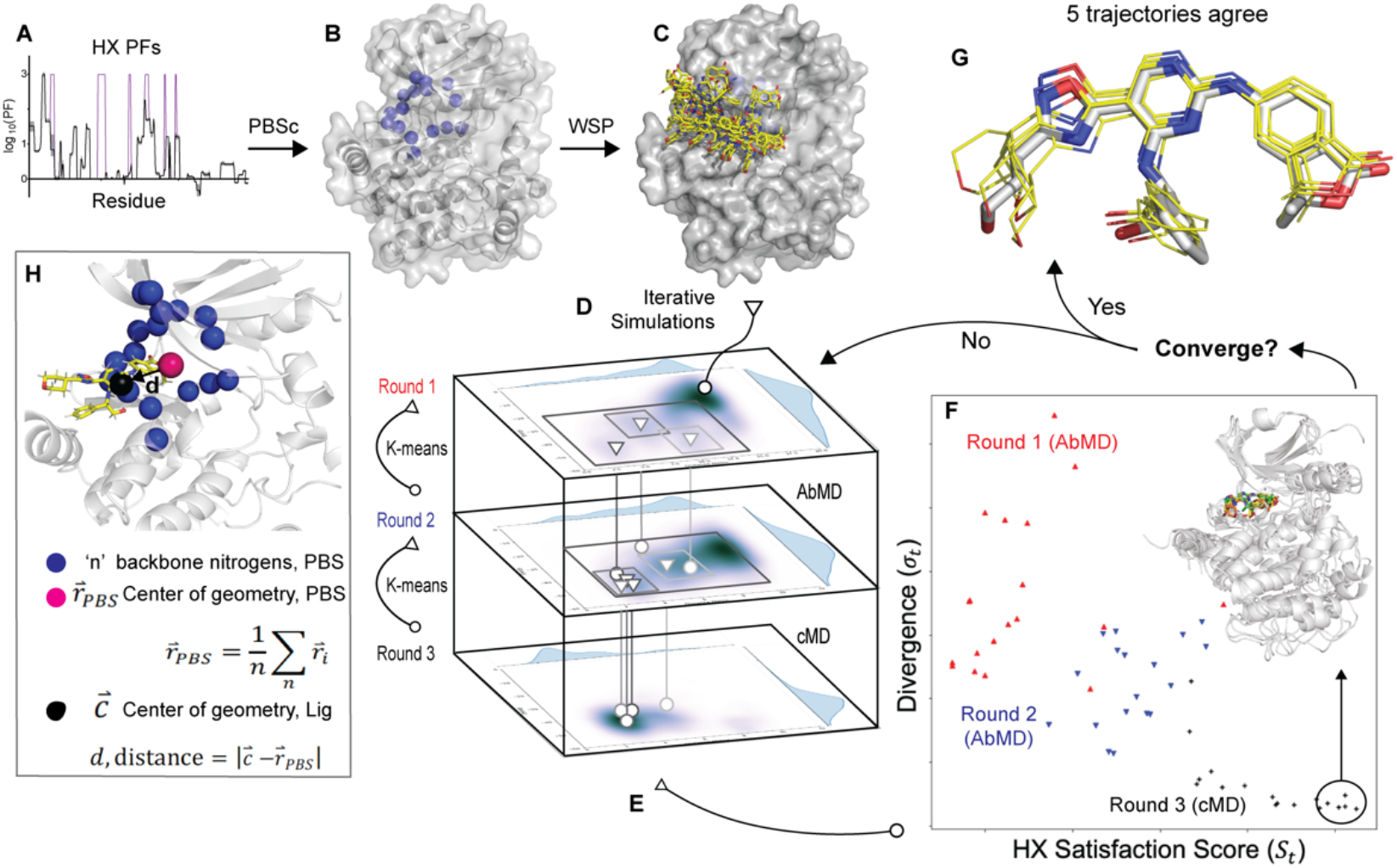
The main components of HX-ESP. **A** PF map and structure submitted to the PBS construction algorithm. **B** Resulting PBS, shown as blue spheres. **C** Starting or seed structures produced by the WSP algorithm, **D** iterative simulations initiated from each seed. All trajectories are combined in each round on the map. The regions submitted for K-means selection is shown in squares in greyscale, matching triangles show the position of the pose selected for the next round. **E** The same K-means selection algorithm is run on individual trajectories to produce a representative structure for scoring. **F** HX and Divergence scores used for convergence testing. **G** Five models that converged are shown in yellow after alignment with the actual bound structure, shown in white. **H** highlights how the PBS is used to create a reaction coordinate or collective variable for simulations. (1.5 column figure).

#### 2. The PBS Construction Algorithm

The perturbation set (PBS), or a list of residues determined from experiment, serves four primary roles in the HX-ESP method. First, this group of atoms defines a region of the protein where the compound is expected to reside, providing a reaction coordinate to monitor progress. This is then used to determine a starting location for initial MD structures (section 3) and allows the automated generation of a collective variable for adaptive-biasing MD simulations. Second, it can be used to confine the compound to a small area of space near its binding site, effectively increasing the compound concentration according to the HX data (section 4). Third, it aids in designing a phenomenological scoring function that combines estimated binding energy from the simulations with the reaction coordinate and plays a role in convergence detection (section 5). Fourth, it provides a subset of residues in the protein for use in clustering applications, such as the K-means algorithm (section 6) used here to select structures from groups.

The PBS construction algorithm (PBSc) takes a PF map, a protein structure or model, and a user-defined threshold PF value as inputs and returns the PBS list (**Figure 1B**). It works by first culling the data, leaving only residues with PF magnitudes exceeding the supplied threshold. Mapping these onto the provided structure, the PBSc then uses a cone-selection algorithm (**Extended data 1.2**) to further cull the list on the basis of their location and orientation in the input structure resulting in the PBS as an output. As the only adjustable parameter, the supplied threshold should include approximately the upper quartile of PF magnitudes. The PBSc therefore objectively defines which residues should be used to guide HX-ESP without relying on subjective interpretation of the HX data.

#### 3. The PBS distance reaction coordinate and starting diversity

The goal in generating seed or starting structures for HX-ESP, is to create a large diversity of ligand orientations as close to the center of the PBS as possible. To begin, we first define a distance between compound and PBS center that will be used throughout HX-ESP as a reaction coordinate (**Figure 1H**):

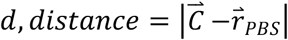

Where the center of PBS, 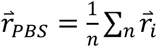, is defined by the position of each amide residue, 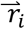, in the PBS, and the center of the ligand, 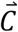, defined by the average position of each heavy atom in the ligand.

For the generation of each starting structure, an algorithm named the Water Shell Populator (WSP, **Figure 1C, Extended Data 1.6**) takes as input the structural coordinates of the protein and ligand, and produces random ligand orientations that minimize *d* and position one ligand atom on the first water shell of the protein. WSP runs until it generates a user-determined number of structures, each with a unique orientation of the ligand. Occasionally modifications to the PBS are needed to generate starting structures using WSP (**Extended Data Figure WSP, Extended Data 1.7, 1.16**). Typically, 20 structures are generated as seeds for the first round of MD, but the number of seed structures can be increased for additional sampling if convergence criteria described in section 5 are not met.

#### 4. Iterative Molecular Dynamics

Iterative swarms of simulations (**Figure 1D**) starting from the structures produced by WSP are used to explore the HX determined binding region, and then to distinguish highly populated states using the PBS. This exploration is initiated with passing seed structures from WSP through two rounds of accelerated sampling that utilize adaptive-biasing MD (AbMD^25,26^, **Extended data 1.10**). Between rounds of MD, a novel k-means selection algorithm (**Extended data 1.8**) is used to cluster highly populated states, and then by using slices of the distance x energy reaction coordinate (**Figure 1H**), generate seed structures for the five replicate trajectories executed in subsequent round. Up to five structures were produced as outputs by this method to seed subsequent simulations evenly with an accurate representation of states observed in previous rounds. Two or more rounds of conventional MD (cMD, **Extended data 1.9**) follow AbMD to relax structures from unnatural forces introduced during accelerated sampling and to continue exploring the pocket. Naturally, as the routine progresses, fewer states will be highly populated, suggesting that the system has started to converge. In the final stage of differentiation, the k-means algorithm is used to select one structure per trajectory, and these structures are compared for convergence based on their HX satisfaction score, detailed below. Convergence is defined as five trajectories producing the same disposition of contacts between protein and ligand. If these criteria are not met, the structures can be resubmitted for another cycle of AbMD and cMD.

Each AbMD round utilizes a center of mass collective variable between the ligand and a positionally restrained atom (referred to as the fiducial atom, **Extended data 1.4**), chosen before each round. Meanwhile, a flat-bottomed harmonic restraint (**Extended data 1.5**) couples 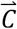 and 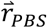, only at the edge of a user-defined distance (typically 8 Å). This minimally affects exploration occurring in the binding region and prevents the compound from being forcibly dissociated by AbMD. The approach accelerates sampling in the region of the molecule defined by experimental data. Up until the final round of cMD, the same flat-bottomed restraint introduced earlier may be used to prevent dissociation. However, in the final round, where output structures will be judged for convergence, this restraint is not used.

#### 5. Convergence Criteria

Individual trajectories are selected for k-means evaluation (section 6) that maximize an HX Satisfaction Score (*S*_*t*_) and minimize Divergence (σ_*t*_):

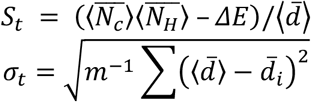

Angled brackets, ⟨***x***⟩, and bar, 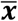, denote the mean value of x either across all m-frames of a trajectory, or over all n-residues in the PBS, respectively. 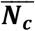represents the mean number of heavy atoms within 6.5 Å of each backbone nitrogen in the PBS; 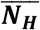, the mean number of H-bond acceptors (oxygen and nitrogen) within 2.5 Å of each backbone nitrogen in the PBS; and **Δ***E* the interaction energy between compound and protein averaged over the trajectory as computed by 1A-MM/GBSA^27,28^.

To be considered convergent (**Figure 1F, G**), five unrestrained sibling cMD trajectories (i.e., trajectories running in the same round) must rank among the highest-scoring trajectories in the round. They must, accordingly, not dissociate over a period of 100 ns, and they must produce structures with minimal deviation between them. Note that more than one pose can be considered converged, and while convergence criteria was met for all test cases considered here, there is no guarantee that convergence criteria will be met.

#### 6. K-means selection algorithm

HX-ESP utilizes K-means for clustering PBS-Ligand related states within/across MD trajectories to identify a set of most representative structure(s) for further analysis **(Figure 1E)**. To determine an appropriate number of clusters to use for partitioning the data, HX-ESP utilizes the Davies-Bouldin index^29^ (DBI), a metric designed to guide unsupervised cluster seeking algorithms. The DBI takes a minimum when clusters are maximally similar by decreasing data feature density as a function of intercluster distance (e.g. the density of data features should be a minimized in between clusters). The number of clusters selected for k-means clustering by HX-ESP corresponds to the first local DBI minimum. The model produced corresponds to the centroid of the most populated cluster when applied to single trajectories in HX-ESP (i.e. for convergence testing). There are exceptions to this rule when producing intermediate structures from simulation swarms that are used to seed downstream simulations (**Figure 1D**) during iterative MD (**Extended data 1.8**). Among the manifold options for clustering MD trajectories^30^, K-means was particularly attractive because the algorithm does not seek to partition data into equally populated clusters and it requires minimal a priori knowledge of the system.

## Results

### HX Experiments

HX protection factor maps for PAK1 and HPK1 show the largest PFs near the binding pocket for all targets once mapped onto the structure (**Figure 2**). Coverage of the sequence was greater than 95% for both HPK1 and PAK1, regions lacking coverage were assigned PF=1. Upon manual inspection of the data, if exchange traces diverged in early or late timepoints (see **Extended Data Figure 2 A-C**), those peptides were flagged and set to the maximum observed PF in that dataset recognizing that the empirical PF could underestimate the true value in this case^31^. The empirical PF was used for all other peptides (see **Extended Data Figure 2 D-F**) yielding PF maps (**Figure 2**) used for PBS construction. These maps were produced after sampling exchange at two different pHs. Only one region of HPK1 was sensitive to pH changes, as determined by the departure from monotonic growth after processing the data as described in the methods (**Extended data 1.11, Extended Data Figure 2C** shows this region in HPK1).

**Figure 2.**
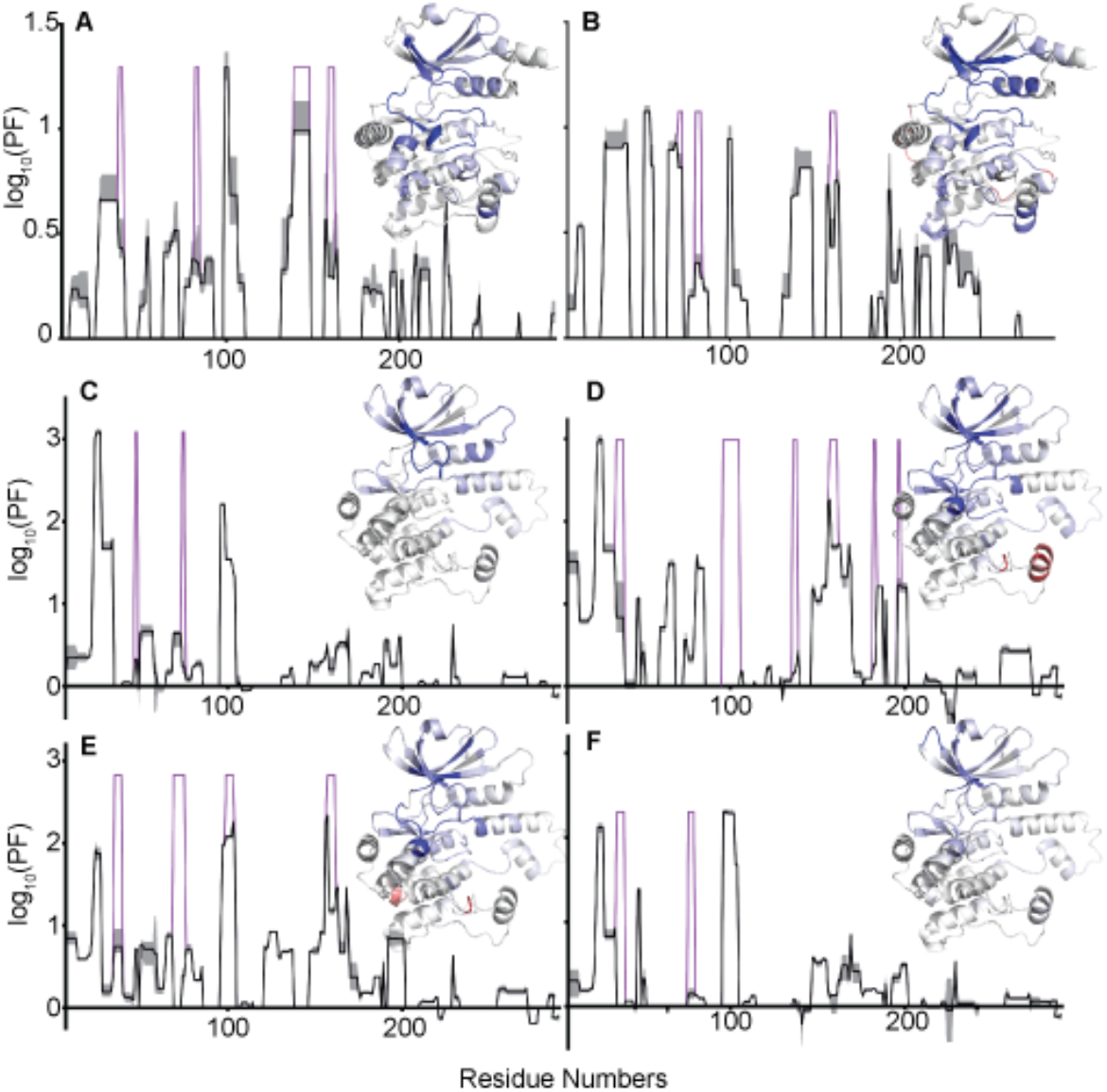
PF plots for all seven ligands. Panels **A** and **B** show PAK1 results for C1 (compound 1), and C2. Panels **C, D, E**, and **F** show HPK1 results for C3, C4, C5, and C6, respectively. Each panel shows empirical PF measurements available for those residues in black with certainty given by the grey shaded region. Magenta lines highlight manually flagged peptides, set to the largest observed PF in that particular dataset. Each panel also shows the PF information colored onto the input structure used for testing HX-ESP. Dark blue shows the maximum log10(PF) value scales linearly to white, where log10(PF)=0. Independently, the minimum log10(PF), showing faster exchange in the bound state, is shaded dark red, scalinng linearly to white, as before. (1 column figure)

### Crystal Structures

Eight novel crystal structures were solved to benchmark the HX-ESP method: three structures of PAK1, wherein limited protein motions are required to accommodate inhibitors, and five of HPK1, where multiple binding modes of inhibitors to different sites required moderate to significant conformational rearrangements. PAK1 and HPK1 structures, conditions, refinement parameters, ligand density maps, etc. are described in **Extended data 1.14-15, Extended Data Tables 4, 5** and **Extended data Figure 3**, respectively. To generate a single monomeric, unliganded structure for each kinase to use as the input structures for HX-ESP, the Type 1 inhibitors (i.e. DFG-D-in, αC-in) in STR3448 and STR5205 (replace with PDBID) were manually deleted for PAK1 and HPK1, respectively, leaving two PAK1 inhibitors (C1, C2 **Figure 3 A,B**) and four HPK1 (C3-6, **Figure 3 C-F**) for testing HX-ESP performance. These two unliganded X-ray structures are referred to as “inputs” and liganded X-ray structures are referred to as “targets” for convenience in subsequent results and discussion sections.

**Figure 3.**
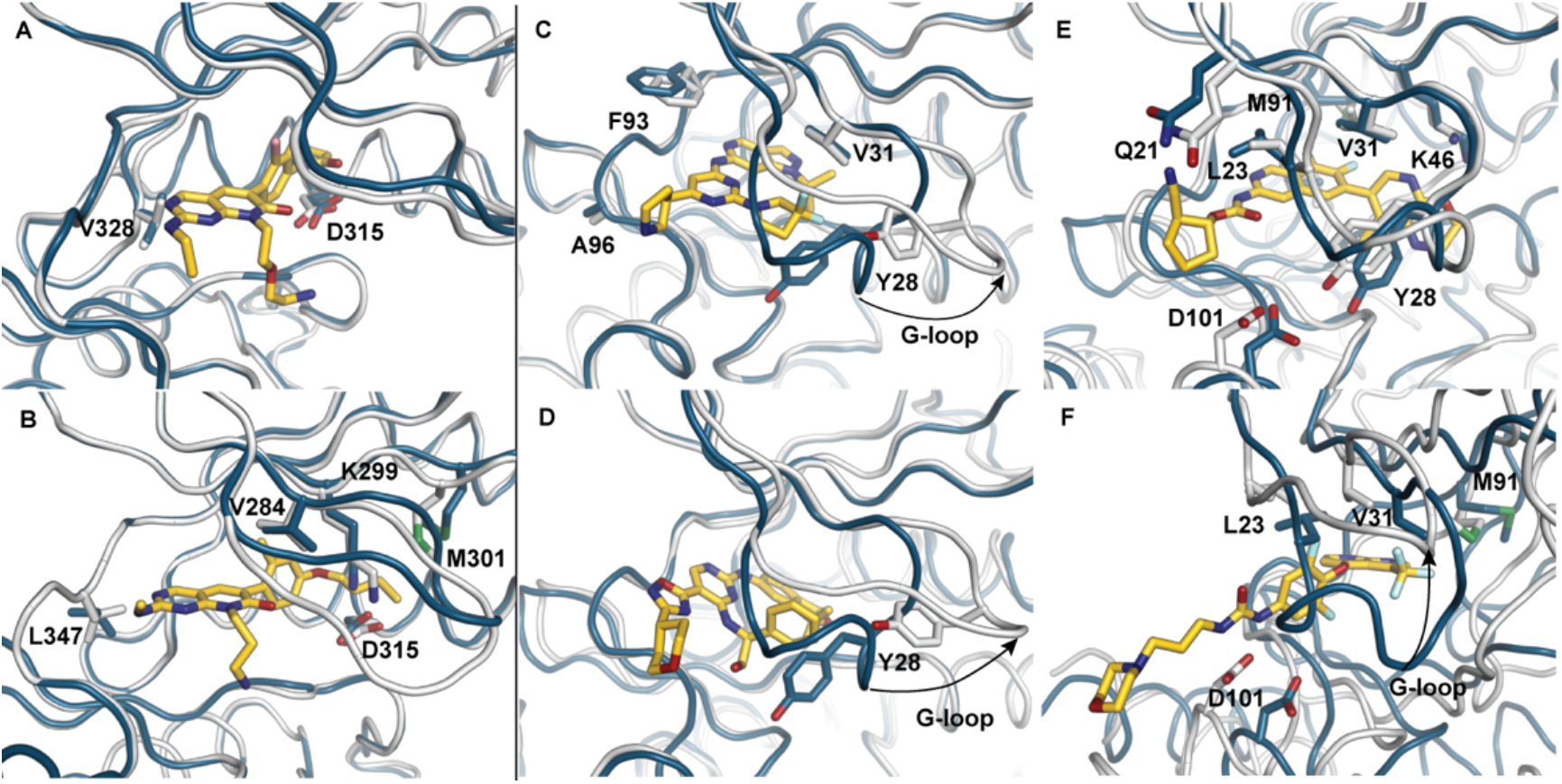
Input (cyan) and target (white) X-ray structures for benchmarking, aligned using atoms within 8 Å of the ligand. Panels A, B show C1 and C2 ligands bound to PAK1, respectively and panels C-F show C3, C4, C5, and C6 ligands to HPK1, respectively; ligands are shown in yellow. (1.5 column figure)

### Evaluation of HX-ESP Performance

The proceeding HX-ESP predictions used the general method described earlier. These models are not ensemble averaged and are selected from a single frame of a trajectory deemed convergent; therefore, we expect deviations to crystal structures on the order of ~1 Å as is commonly seen in simulations and recently validated to exist in solution by NMR^32^. Further, because all crystal structures show domain swapped dimers, deviations from crystal structures in HX-ESP models are expected due to refolding of these domain swapped elements. To account for these differences, three X-ray relaxed models were prepared from each target crystal structure (**Extended data 1.17**). These X-ray relaxed models establish a baseline, as contacts lost due to refolding of domain-swapped elements are also expected to be absent in HX-ESP models. For the purpose of benchmarking against currently available software, binding modes were also predicted using the data-driven, flexible docking program HADDOCK (version 2.4) ^33–35^.

HADDOCK was guided by ambiguous interaction restraints (AIRs) determined by the PBSc algorithm developed for HX-ESP and all other settings followed recommendations from the developers for small molecule docking. HADDOCK’s best scoring cluster of four models was taken as the flexible docking result for each target. Thus, for each test ligand, five convergent HX-ESP models, four flexible-docking models, and three X-ray relaxed models are produced. Five metrics were used to assess similarity to target structures: (1) backbone RMSD in the vicinity of the binding pocket (**Table 1**), (2) mean, all-atom ligand RMSD after pocket alignment (**Table 1**), (3) interatomic ligand-protein contact distance differences between models and the target X-ray structure (**Figure 4**), (4) the mean and standard deviation over all models using a contact distance difference RMSD computed across all models (**Extended Data Table 1**), and (5) the absolute number of interatomic contacts between the ligand and target as observed in target X-ray structures. To define the presence of a particular contact, a max distance of 3.5 Å for H-bonds and 4.5 Å for non-polar interactions was imposed and a contact is considered to be correctly predicted if observed within these limits for more than 50% of models produced by the technique (**Extended Data 1.18, Extended Data Tables 2, 3**). Successful predictions should have pocket, ligand, and contact RMSDs less than 2 Å, a value commonly used to evaluate docking performance^36^, and identify contacts observed in reference X-ray structures, particularly those contacts to the kinase hinge that are observed in all known kinase inhibitors. See **Extended data 1.13** for additional information.

**Table 1.**
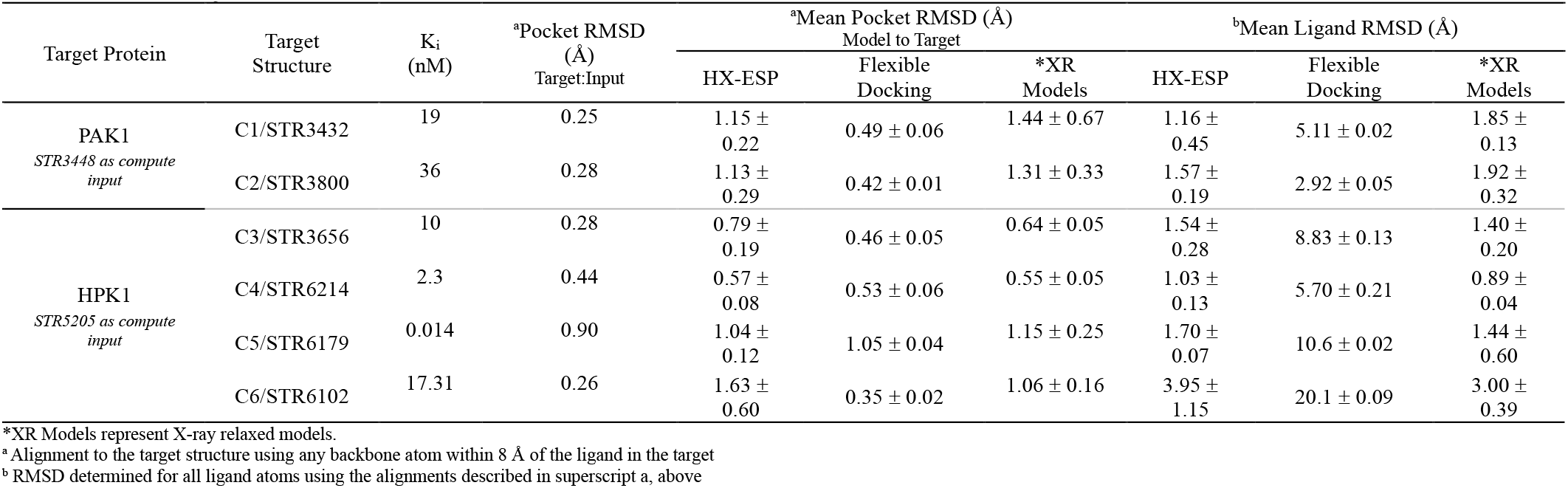
Summary of results.

**Figure 4.**
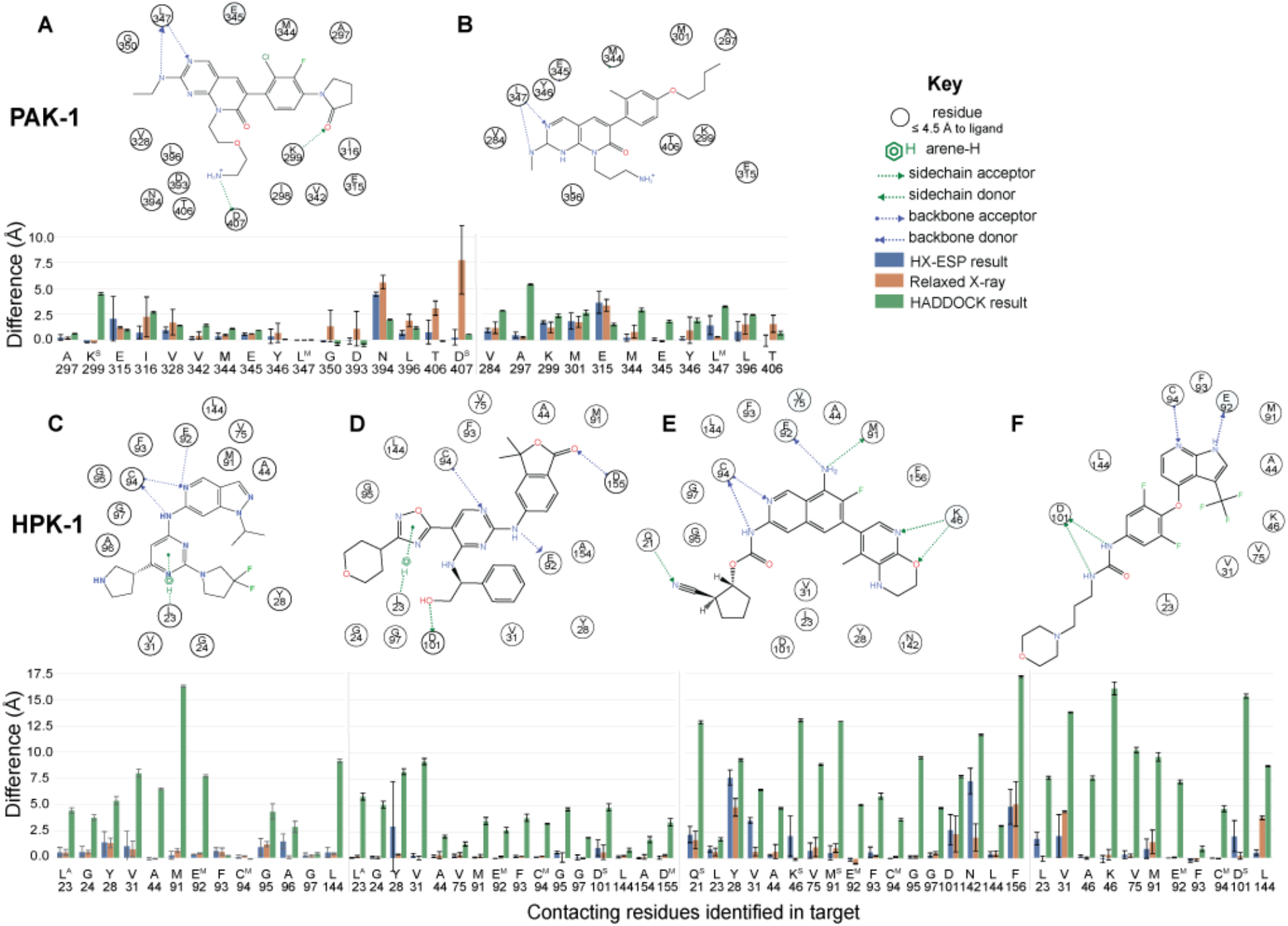
Mean ± Std.Dev. of contact distance differences between contacts found in bound X-ray structure and the distance between equivalent atoms in models shown for flexible docking (green, n=5), relaxed X-ray (orange, n=3) and HX-ESP (blue, n=5). Panels A and B show PAK1 results for C1 and C2. Panels C-F show HPK1 results for C3, C4, C5, and C6, respectively. Superscripts M, S, and A highlight main chain H-bonds, side chain H-bonds, and Arene-H interactions, respectively. The mean over all contacts are given for residues that contact multiple ligand atoms. Diagrams of each contact, produced by MOE in 2D, are included above each bar plot. (2 column figure)

### PAK1 Benchmarking

PAK1 targets are similar to the input structure having pocket backbone RMSDs of ~0.3Å (**Table 1**) with respect to the input structure used for computational predictions. The binding pocket is well formed in the PAK1 input structure and can bind all test ligands with minor conformational change making these ligands easier to predict. All test ligands bind with the same type-I mode of inhibition (**Extended Data 1.14.1**). Overall, despite flexible docking achieving lower pocket RMSDs (**Table 1**), HX-ESP demonstrated superior accuracy in capturing the binding interactions for both PAK1-C1 and PAK1-C2 complexes.

#### PAK1 bound to C1

***(See Figure 3A, Figure 4A, Figure 5E, Table 1, Ext. Data Tables 1-3)***

**Figure 5.**
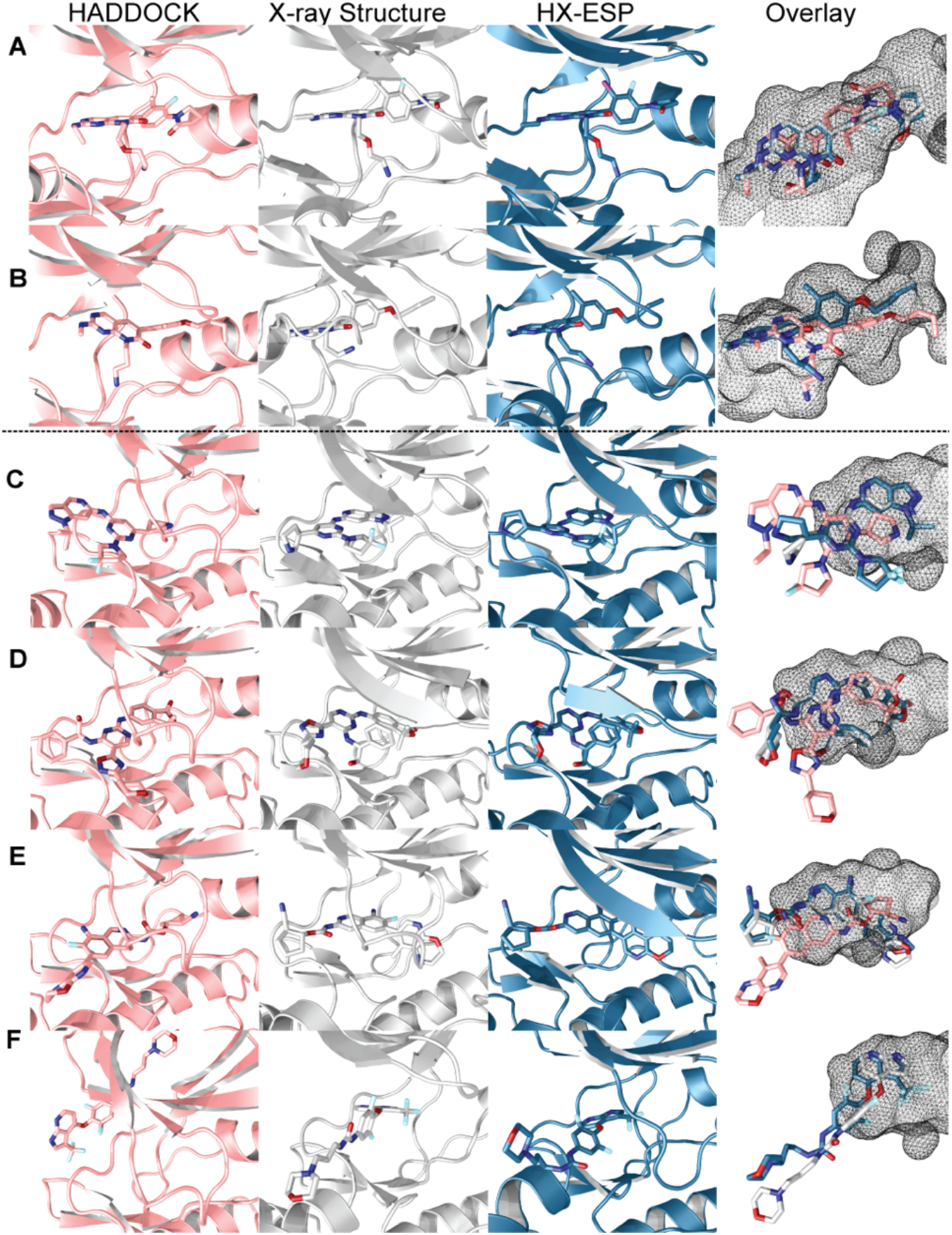
Models are aligned to the unbound input structure using the same residues used to compute mean pocket RMSD (Table 1). Panels A and B show PAK1 results for C1 and C2, respectively. Panels C-F show HPK1 results for C3, C4, C5, and C6, respectively. Each panel has four insets showing the flexible docking (pink), the target bound X-ray structure (white), and HX-ESP (blue). The cartoon is stripped away in the fourth inset for each panel and the binding pocket as observed in the input structure shown in mesh to illustrate how the ligand is positioned by HX-ESP and flexible docking relative to the target structure for each ligand.

HX-ESP models have a mean pocket RMSD of 1.15 Å, similar in magnitude to the 1.44 Å observed in X-ray relaxed models, with a mean ligand RMSD of 1.16 Å. These models reproduce 13/16 contacts observed in the target structure with a mean contact RMSD of 0.98 Å. All H-bonds are identified with the exception of a sidechain H-bond between the D407 and ligand terminal amine which exceeds distance thresholds in one model. Notably, the electron density was weak for this terminal amine (**Extended data Figure 3A.C**). V328 oscillates away from and towards the ligand due to the orientation of the backbone exceeding the distance threshold in every model. A polar interaction seen in the target structure with E315 is observed in 40% of models. All contacts missed by HX-ESP are also lost during the relaxation of the X-ray structure where only 9/16 contacts were preserved.

Flexible docking models give a pocket RMSD of 0.46 Å, and contact RMSD of 1.16 Å; however, they exceed the 2 Å threshold for ligand RMSD. We consider these predictions successful because they find a total of 10/16 native contacts, including the critical backbone hinge interactions with L347 and E345. Due to a 180° rotation of the pyrrolidone, multiple contacts are missed; however, the compound occupies the same space in the binding pocket. While HX-ESP reproduces the binding mode better than flexible-docking, both techniques perform well for this ligand.

#### PAK1 bound to C2

***(See Figure 3B, Figure 4B, Figure 5F, Table 1, Ext. Data Tables 1-3)***

HX-ESP models have a mean pocket RMSD of 1.13 Å, similar in magnitude to the 1.31 Å observed in X-ray relaxed models with mean ligand and contact RMSDs of 1.57, and 1.25 Å, respectively. Closer inspection of each contact shows that 6/11 contacts are identified. While the critical hinge H-bond to L347 lies just beyond the 3.5A threshold on average, HX-ESP models satisfy this contact with a neighboring nitrogen due to being slightly shifted further into the pocket than observed by X-ray models. This shift also causes a loss of the contact to V284. A rotation of the butyloxy tail places K299, M301, and E315 just beyond the 4.5 A threshold. In total, 6/11 contacts are properly formed, the compound sits in roughly the same position observed in the target structure and while a register shift results in a different H-bond to the hinge, the importance of engaging L347 remains evident in these models.

Flexible docking models give a pocket RMSD of 0.42 Å but exceed the 2 Å threshold for both ligand and contact RMSDs with values of 2.92 and 2.55 Å, respectively. Though only 2/11 total contacts (both non-polar) are identified, the ligand is placed in a roughly correct orientation. The majority of missed contacts were only 1-2 Å beyond the contact threshold and many would be expected to form if these models were allowed to relax freely using MD simulations similar to those used to relax the X-ray models. All-in-all, HX-ESP outperformed flexible docking for C2.

### HPK1 Benchmarking

The binding pocket volume in the input X-ray structure must expand by more than 250% to properly engage any of the test ligands (see **Extended Data Figure 4**) because the bound ligand that was deleted to produce an unliganded input structure for calculations is much smaller than any of the ligands tested here. In liganded structures for C3, C4, and C6, the pocket opens by extending a beta sheet that comprises G-loop residues 23-25, 30, and 31 (**Figure 3C, D, F**) resulting in backbone displacements that exceed 8 Å. In the case of C5, the G-loop must move by a similar magnitude to provide access to the pocket, but can adopt a conformation similar to the input structure once bound (**Figure 3E**). Therefore, binding of each test ligand will require large backbone displacements in the input structure making these predictions much more difficult than PAK1. Further, unlike PAK1, HPK1 ligands span multiple modes of inhibition (**Extended Data 1.14.2**). For HPK-1, HX-ESP models achieved a mean ligand RMSD of 1.5 Å over all compounds, accurately capturing the binding interactions and identifying crucial contacts. In stark contrast, flexible docking exceeded the 2 Å threshold for ligand RMSD in every case, failing to produce reliable predictions for any of the test ligands, underscoring HX-ESP’s superior accuracy and reliability in cases where a binding pocket must significantly change in order to bind.

#### HPK1 bound to C3

***(See Figure 3C, Figure 4C, Figure 5C, Table 1, Ext. Data Tables 1-3)***

HX-ESP models capture the binding pose correctly with a mean pocket and ligand RMSD of 0.79 and 1.54 Å, respectively, and a mean contact RMSD of 0.86 Å. These models reproduce 11/13 total contacts, including the hinge H-bonds to E92 and C94. Of the two missed contacts to G95 and A96, both are present in two of the five converged results. Both residues are roughly in the same conformation as the target structure but lie just beyond the contact threshold in cases where they are missed. Alternatively, while flexible docking models give a pocket RMSD of 0.46 Å to the target structure, these models do not produce the correct bound pose, though they do contain the critical hinge H-bond to C94 along with two other contacts observed in the target structure.

#### HPK1 bound to C4

***(See Figure 3D, Figure 4D, Figure 5D, Table 1, Ext. Data Tables 1-3)***

HX-ESP models have a mean pocket RMSD of 0.57 Å, similar to the 0.55 Å observed in X-ray relaxed models with mean ligand and contact RMSDs of 1.03, and 0.58 Å, respectively. Nearly all contacts are identified by HX-ESP, including all backbone and sidechain H-bonds. 15/16 contacts are correctly identified lacking only the contact to Y28, a residue on the highly dynamic G-loop. This residue takes different conformations in all five converged models, agreeing with the target structure in only one. Despite the large number of conformational differences between input and predicted models, nearly every sidechain takes a similar conformation to the target X-ray structure and HX-ESP models accurately reproduce the binding pose for C4. Flexible docking failed to reproduce the correct binding pose for HPK1 bound to C4.

#### HPK1 bound to C5

***(See Figure 3E, Figure 4E, Figure 5E, Table 1, Ext. Data Tables 1-3)***

HX-ESP models have a mean pocket RMSD of 1.04 Å, similar to the 1.15 Å observed in X-ray relaxed models with mean ligand and contact RMSDs of 1.70, and 2.17 Å, respectively. Critical contacts to the hinge residues were observed in all HX-ESP models, but the apical pyrido-oxazine moiety is flipped by roughly 180 degrees compared to the target structure, disrupting contacts to Y28, V31, K46, and N142 by roughly 2.5 Å. Though this apical arm was flipped, it was placed in the same space. This resulted in finding only 8/17 contacts in HX-ESP models compared to the 10/17 preserved in X-ray relaxed models where no flipping occurred. Excluding contacts involving the pyrido-oxazine, HX-ESP models correctly identify all other contacts, including the hinge H-bond interactions, and thus largely succeed in producing the correct binding mode. C5 was unique among ligands tested because convergence criteria were not initially met. Increasing the number of starting structures from 20 to 40 (see section 3) was required to converge and produce the result shown here. In flexible docking models, the compound is flipped along its long axis and does not access the back pocket; none of the 17 contacts present in the target structure are correctly identified.

Following up on the flipped pyrido-oxazine moiety revealed that HX-ESP never sampled the correct pose of this ring; had it been sampled, it would have been ultimately selected as the preferred structure on the basis of X-ray relaxed simulations showing a higher satisfaction score for the correct pose. Notably, comparison of the bound structure (PDBID: XXX) to the input structure used for these simulations (PDBID: XXX) reveals that the domain swapped structures have different folds in residues that contact the ligand. We believe HX-ESP could not sample the correct pose for this ring due to refolding of the critical, domain-swapped contacts needed for stabilizing that orientation having not reached the proper configuration over the course of simulations.

#### HPK1 bound to C6

***(See Figure 3F, Figure 4F, Figure 5F, Table 1, Ext. Data Tables 1-3)***

HX-ESP models have a mean pocket RMSD of 1.63 Å, similar to the 1.06 Å observed in X-ray relaxed models with mean ligand and contact RMSDs of 3.95, and 1.01 Å, respectively. The anchoring trifluoromethyl dihydro pyrrolopyridine group overlays with the target structure with very little deviation (**Figure 5D**), and all hinge H-bonds are correctly formed. Of the three missed contacts, all involve the solvent exposed tail which takes a different pose in each of the five converged models, consistent with the weak electron density observed for this portion of the compound in the target liganded structure (Extended data Figure 3B.d). HX-ESP finds 9/11 contacts observed in the target structure, including the two critical backbone H-bonds to hinge residues E92 and C94. Despite the heterogeneity of tail configurations that explain missed contacts to L23 and V31, every model finds an H-bond to the D101 sidechain, only two of the five models contain the H-bond observed in the target structure. Despite missing the contact by thresholds set for this analysis, these models do highlight the importance of interacting with D101 and generally capture the correct binding mode for C6. Perhaps the often used 2 Å ligand RMSD threshold^36^, was too strict for HPK1 where a roughly 2 Å ligand RMSD exists between three different sunitinib-bound HPK1 structures^37^, highlighting the structural plasticity of this kinase. Flexible docking models place the compound outside of the binding pocket, failing by all metrics to predict the bound pose of C6.

## Discussion

In this study, we evaluate the effectiveness of Hydrogen Exchange (HX) data for predicting ligand-protein interfaces through HX driven *E*xperimental *S*tructure *P*rediction (HX-ESP). HX-ESP utilizes a combination of in-vitro measurements and in-silico calculations to predict and validate binding modes of ligand-protein systems. HX data, extracted in the form of Protection Factors (PFs), defines how strongly a ligand effects the dynamics of the protein across each amino acid. The PFs, are then turned into a relevant set of amino acids, called the perturbation set (PBS). The PBS is utilized throughout the in-silico component for constructing collective variables for accelerated simulations, and for targeted analysis of extracted simulation data. By running iterative swarms of accelerated then standard molecular dynamics, we demonstrate that the binding mode for static and dynamic ligand-protein systems can be accurately predicted.

We chose to benchmark the method against kinases due to the well-studied ATP binding pocket and known contacts to the kinase hinge critical for affinity^38,39^ providing a robust basis for performance evaluation, particularly with the additional challenge of the large conformational changes in HPK1. Two kinases, PAK1 and HPK1, were selected as targets for blindly testing the HX-ESP routine. PAK1 was chosen because it has a relatively open binding pocket in a majority of structures, requiring only a small reorientation of residues that contact the ligand regardless of ligand size. Conversely, HPK1 has a much more dynamic binding pocket that collapses in the absence of ligand. None of the tested ligands fit without extensive conformational change making it much more difficult to correctly predict the binding mode than PAK1.

The results from our benchmarking finds that in every case, HX-ESP correctly identifies critical hinge contacts and the majority of non-polar contacts found in target structures, despite the deviations that occur in HX-ESP simulations as a result of refolding domain swapped elements. The majority of contacts missed by HX-ESP were also lost by X-ray relaxation models, suggesting either that they are less important for affinity, or simply lost due to refolding of domain swapped elements. However, there were a few notable exceptions. Consider the H-bond interactions K46 (HPK1+C5), D101 (HPK1+C6), and L347 (PAK1+C2) that were preserved during X-ray relaxation but not identified by HX-ESP. In every case, one or more models in the HX-ESP converged ensemble identified the contact (**Figure 4, Extended Data Table 2**). This remained true for non-polar contacts as well. Thus, using mean contact distance differences, or requiring more than 50% of models to possess contact distances below a certain threshold clearly underestimates the value of HX-ESP akin to the difference between mean and mode - the average does not always provide a representation of reality.

### The advantage of HX-ESP

The success of HX-ESP depends most heavily on the experimental information that drives the algorithm. Input data from HX experiments work so well because the extracted PF (protection factor) proportionally estimates how much work has been done locally on the protein upon binding. The relative magnitude of a PF must inversely relate to its proximity to affinity-driving interactions due to energy dissipation. The PBSc algorithm introduced here uses this principle to unambiguously determine which residues most likely interact with the ligand when provided an unliganded input structure. Where sorting HX changes into direct and indirect effects of binding was variably left to individual interpretation, the PBSc algorithm provides a reproducible method that works for the first time to our knowledge.

To calibrate the results of HX-ESP, a popular data-driven docking program called HADDOCK was used to provide a flexible docking benchmark. HADDOCK was chosen because it could use the HX-ESP PBS defined by the PBSc algorithm to define AIRs (Ambiguous Interaction Restraints). Furthermore, HADDOCK utilizes iterative rounds of coarse-grained MD, and it even considers solvent interactions^34^, making it one of the most advanced docking solutions available. The PBSc algorithm eliminated ambiguity in selecting AIRs guided HADDOCK to the correct answers in two of three test cases against PAK-1. HADDOCK was able to identify critical hinge contacts to HPK1 with C3; however, due to restriction of motion, flexible docking was unable to correctly predict any of the compounds bound to HPK1. Flexible docking algorithms, like HADDOCK, are most attractive in situations where the binding pocket is well-formed in an unliganded state and may even be preferred due to low computational cost.

While adaptive-biasing proved to be a very useful component of HX-ESP, other biasing or accelerated methods should be investigated in lieu of it. One of the major drawbacks of adaptive biasing is the requirement of knowing the end state. HX-ESP overcomes this by creating a diversity of ligand starting positions, and progressing several swarms of simulations through to convergence. While this helps resolve the weakness of adaptive biasing, there may be other approaches more cost efficient, like supervised-MD^40^, or weighted ensembles^41^.

We have shown that HX-ESP is a powerful method for structural prediction of small molecules bound to protein targets. It reliably found the correct binding pose even when the binding pocket in HPK1 was less than half the volume needed for inhibitor binding, suggesting a potential utility against targets where the binding pocket has not formed. Future work will focus on the ability of HX-ESP to predict lower affinity ligands (i.e. fragments) to guide rational design in the absence of experimental structures.

## Code and structure availability

All structures referred to in this manuscript are deposited in the RCSB and PDBIDs are matched with names used in here in **Extended data 1.14**. HX-ESP was developed in parallel to structure refinement and used structural coordinates prior to a final round of refinement to seed X-ray relaxed simulations, provide input structures to HX-ESP, and as targets for performance analysis. Input (Apo), target (bound), and all models produced as described in this manuscript and used to compare accuracy are currently being uploaded to an online repository and will be made available.

## Author Contributions

BW and AP conceived of HX-ESP, performed simulations, and analyzed the results. BW performed and analyzed HX experiments. WW picked compounds for benchmarking (BW and AP were blinded to their binding modes). WW and PW oversaw generation of crystals and initial refinements, JK further refined structures for upload to the RCSB. BW, AP, and JK wrote the manuscript.

## Acknowledgements

The research was funded entirely by Roche, Inc. through the Genentech gRED Incubator program. We thank Joachim Rudolph and Bryan Chan who provided compound synthesis information and were instrumental in releasing the supporting data package for this work, and Peter Hsu, who graciously refined PAK-1 structures reported here. We thank Melinda Mulvihill for critically reviewing the manuscript. We thank our sponsors, Wayne Fairbrother, and Eric Fahs and his team for their support and strategic planning, without whom this work would not have been possible. We thank the groups at ChemPartner and Viva Biotech for contributions to the crystal structures. Data were collected at Southeast Regional Collaborative Access Team (SER-CAT) 22-ID-D (or 22-ID-E) beamline at the Advanced Photon Source, Argonne National Laboratory. SER-CAT is supported by its member institutions, equipment grants (S10_RR25528, S10_RR028976 and S10_OD027000) from the National Institutes of Health, and funding from the Georgia Research Alliance. This research used resources of the Advanced Photon Source, a U.S. Department of Energy (DOE) Office of Science user facility operated for the DOE Office of Science by Argonne National Laboratory under Contract No. DE-AC02-06CH11357. Beamline 8.3.1 at the Advanced Light Source is operated by the University of California at San Francisco with generous grants from the National Institutes of Health (R01 GM124149 for technology development and P30 GM124169 for beamline operations), and the Integrated Diffraction Analysis Technologies program of the US Department of Energy Office of Biological and Environmental Research. The Advanced Light Source (Berkeley, CA) is a national user facility operated by Lawrence Berkeley National Laboratory on behalf of the US Department of Energy under contract number DE-AC02-05CH11231, Office of Basic Energy Sciences. We thank the Shanghai Synchrotron Radiation Facility (SSRF) and staff of beamline BL18U1 for crystallographic data collection.

