## SupplementaryInformation for "Integrating Hydrogen Exchange with Molecular Dynamics for Improved Ligand Binding Predictions"

January 13, 2025

#### 1 Extended Data

##### 1.1 General Definitions

Terms used throughout these extended methods are defined here for convenience.

The PBS (perturbation set) is a collection of $n$ atoms, defined from the data and cone selection algorithm described in section 1.3, PBS construction. Mathematically, the $i^{th}$ residue in the PBS is specified by the position of its amide nitrogen, $\vec{r}_i$, and the center of the PBS:

$$\vec{r}_{pbs} = \frac{1}{n}\sum_{i=1}^{n}\vec{r}_i$$

The ligand has a center of geometry defined by its heavy atoms and is represented by the vector $\vec{C}$.

##### 1.2 Conic Containment Determination

Several techniques within this paper require us to check if an atom lies within a conic volume. Here we outline the process for this in the following procedure

First describe the desired conic volume
$P_0$: Position of cone vertex
$h_0$: distance from $P_0$ to define the start of the volume
$h_f$: final distance from $P_0$ to define the end of the volume
$s$: cone theta
$v$ : unit vector that describes the central axis direction of the conic volume

```
d_cone: Vector3 = v;
dangle: float = cos(theta);
accepted_atoms: list;
while cur_atom_id < total_atoms do
    pos: Vector3 = get_xyz(cur_atom_id) ;
    dpos_unit: float = make_unit_vector(pos - P0)
    dpos_angle: float = dot(dpos_unit,d_cone)
    if dpos_angle ≥ dangle and distance(pos − P0) within [h0, hf] then
        accepted_atoms.append(cur_atom_id);
    end
    cur_atom_id++;
end
return accepted_atoms;
```

The resulting output will be the number of atoms that lie within the defined region.

#### 1.3 PBS construction and Cone-selection

PBS selection begins by thresholding the PF values measured in the experiment. Typically, a value that is roughly 0.8 of the maximum observed PF is chosen (See Supplemental Roadmap for information on the exact thresholds used for compounds in this manuscript). All residues above that threshold are accepted into the initial PBS and then filtered using Conic Containment Determination (1.2) to eliminate residues from the PBS which aren't directly accessible from a compound residing at the center of the PBS (i.e. $\left|\vec{C} - \vec{r}_{pbs}\right| = 0$), and then removed from the PBS as they cannot favorably contribute to pocket exploration in the algorithms described here.

The Cone-selection algorithm uses conic containment determination for each residue to determine the identity of any atom in between each residue in the PBS and $\vec{r}_{pbs}$ - parameterized as follows:

$$\vec{P}_0 = \vec{r}_i$$

$$h_0 = 8$$

$$h_f = 15$$

The slope is usually set to roughly 15° but may need to be adjusted for a different target. Once this has been done for each residue in the initial set, only residues with a clear line of sight to the PBS center are kept as all residues with non-neighboring atoms inside their conic volumes have been removed.

This procedure defines exactly how the PBS is constructed from measurements.

#### 1.4 Fiducial Atom

One of the challenges faced in this work was protein denaturation when we attempted to use the PBS atoms and the compound atoms as respective masks for a center-of-mass collective variable. To avoid this problem, we chose to use a positionally restrained fiducial atom and the compound for masks to define our adaptive biasing collective variable. The fiducial atom was defined on-the-fly by randomly selecting an atom from a conic volume defined using the direction vector between the ligand and the PBS for each input structure.

The detailed method is as follows: Construct a direction vector between the ligand and PBS center-of-mass. Parameterize the cone selection algorithm with said direction vector with apex at the PBS center of mass. Then choose, at random, an atom that lies 8 Å away from the PBS center of mass and lies within the conic volume. Finally, apply a 100 kcal/mol positional restraint on that atom.

#### 1.5 Flat-bottom restraint

To encourage sampling of a specified region near the PBS center we employ flat-bottom distance restraints to restrict the ligand to a space by the PBS center. In classical molecular dynamics (cMD) simulations, this can be useful to prevent dissociation and increase sampling at higher temperatures but it is required when using AbMD as our strategy using a fiducial atom would immediately result in dissociation otherwise. This flat bottom restraint was implemented in amber using the nft package setting r1, r2, rk2 to 0, rk3 to 100, and finally r3 to $d_0$ and r4 to $d_0 + 1$, where $d_0$ is the desired maximum distance.

#### 1.6 WSP

The Water Shell Populator (WSP) algorithm creates a diverse set of the ligand protein starting structures such that the ligand's rotational and translational conformations are unique, and their distances to $\vec{r}_{pbs}$ are minimized. WSP works by constructing a surface 3A away from the protein utilizing a binary array. The simulation box is filled with empty cells indexed by their position in space. Then, the volume is determined through iterating across all the atoms and enabling the cells that lie with 3A radius of the atom. With the volume of the protein determined, the surface is selected by identifying all false values that neighbor a true value in space. Furthermore, we reduce the surface to a region that lies within a distance from the PBS, most often less than 9 angstroms. With the selected surface region, WSP will randomly populate the a static ligand structure positionally and rotationally onto the protein's surface. Any pose that does not contain atomic clashes is stored for later retrieval. This process repeats until the desired number of structures have been achieved. The resulting distribution of structures is culled to guarantee a minimum RMSD between any two structures in the distribution, such that no two structures are very similar. Finally, we sort the structures by distance to $\vec{r}_{pbs}$ and output the number of requested starting structures.

##### 1.7 WSP Selective PBS Modification

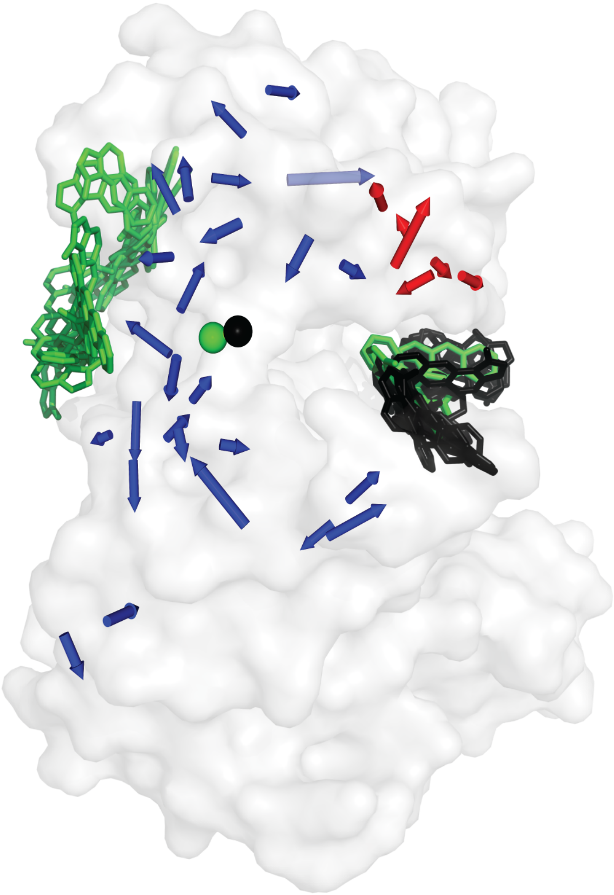

Figure 1: Blue and red arrows take the amide of a particular residue as origin and point towards the furthest sidechain carbon. Using the PBS construction algorithm, blue arrows are selected causing the WSP algorithm to populate starting structures in green. Temporarily adding residues highlighted by the red arrows to the PBS results in the black starting structures, and better WSP performance. Those residues are then removed for downstream use.

Due to the simplistic nature of WSP, sometimes we may need to modify the PBS input for WSP since structures can be populated in regions nearby to the PBS but not accessible to the PBS. As an illustration, see **Extended data figure 1** showing the WSP output for a PAK1 compound using only the PBSc (represented by blue arrows, see PBS, 1.16.1) which results in starting structures shown in green. Looking at the structure reveals that due to the PFs being largest on the back wall of the pocket, compounds are populated on the wrong side of the protein. In a case like this, adding sham residues into the PBS (shown by red arrow, see WSP PBS, 1.16.1) shifts the PBS COM enough to produce the black starting poses. Using the green starting poses does eventually converge to the correct answer, but this easy to do manual intervention allows HX-ESP to converge much faster. The automatically selected PBS by the PBSc, along with the manually adjusted PBS for WSP in one case are given in section 1.16.

Additional changes to the WSP algorithm could be made to avoid this, such as constructing a matrix of distances within the PBS and determining where the largest combination of PBS distances is LEQ to the longest length of the ligand.

#### 1.8 K-means

Clustering is utilized in two separate forms within the analysis of trajectories. The first is in the specific slicing of frames corresponding to a coordinate region and the second is the entire trajectory. In both of these forms, we determine an appropriate number of centroids by utilizing a population number selection protocol.

To select the number of populations, we iterate through up to 20 populations and compute the Davies-Bouldin index [1] (DBI) for each one. The DBI measures how well a clustering routine has maximized the distance between each cluster while minimizing the spread within each cluster. The best number of populations for clustering is determined by selecting the first local minimum in the series of DBI values calculated across 1-20 expected populations.

Clustering is always performed on the position of the ligand and the positions of the PBS residues. In the full trajectory clustering algorithm, we utilize the cpptraj cluster command with kmeans, randompoint enabled, random sieve over 10 frames, and RMS set to target the PBS and compound.
In the slicing protocol, we utilize kmeans within the pytraj package and opted not to change the defaults defined in pytraj.

Full trajectory clustering is utilized for the output structure associated with the HDX score, whereas the trajectory slice clustering is utilized for seeding the next round of sampling in exploratory rounds. When clustering a sliced trajectory, the goal is to select relevant and diverse structures to seed the next round of sampling. If there is one dominant structure, in terms of population, then that structure is selected. However, if the populations of other structures are comparable to the largest populated structure, those can also be progressed. When comparable populations exist, they are manually inspected to determine if the structure is functionally similar to other comparable populations. If true, then only one of the similar structures is chosen to be progressed.

When the goal is to select a single structure from a clustering result, the most populated centroid is selected unless the populations of the second and third most populated centroid are greater in sum than the dominant population. In these cases, the second most populated cluster may be chosen if its trajectory RMSD is smaller than the most populated group. This is done at convergence testing.

#### 1.9 MD Force-field Generation for Ligands

MOL2 files are prepared from chemical SMILES of each compound using OpenEye OEChem TK 2022.2.1 (OpenEye, Cadence Molecular Sciences, Santa Fe, NM. http://www.eyesopen.com) and then geometries optimized prior to partial charge assignment using Gaussian 2009, both completed at the HF/6-31g* level of theory [2] and Antechamber (Amber 2022 [3]). For charge assignment, 10 concentric layers of points are used with roughly 2500 points per atom; sander and tleap (part of the AMBER suite) are then used to prepare force fields for use in AMBER.

All molecular dynamics simulations are performed in the AMBER (v2020 [3]) suite using AmberFF14SB/TIP3P [4] force fields for the target protein. Hydrogen Mass Repartitioning and SHAKE are enabled with a times tep of 0.004 ps. All PAK1 compounds were run at 350K and all the HPK compounds were run at 375K using the constant temperature Langevin Thermostat. The collision frequency of the chosen thermostat was defined to be 3 and utilized a bath coupling constant of 0.5. Constant pressure of 1.0 atm was applied using isotropic position scaling via Berendsen barostat. NFT calculations like distance restraints and AbMD were enabled on runs that contained biasing techniques. All other settings following the defaults listed in the Amber manuals [3]. All simulations were initially minimized using a decreasing positional restraint in conjunction with the AMBER minimizer (imin=1) at intervals of 50kcal, 25kcal, 20kcal, 15kcal, 10kcal, 5kcal with ncyc=100 and mcyc=200. The final minimization run is done with no positional restraint and ncyc=1000, maxcyc=10000. The output of the final minimization round is fed into an iterative round of equilibration to bring the system up to the desired temperature. The first round from 20k to 150k, setting nstlim=5000 and dt=0.001 with solute restrained. The second round from 150k to desired temperature, setting nstlim=50000 and dt=0.001. Then keeping constant temperature and setting nstlim=70000 and dt=0.0005. The final equilibration frame is fed into production for simulations.

#### 1.10 AbMD

The adaptive biasing module in AMBER utilizes metadynamics (flooding) of the energy potential(s) across a specified reaction coordinate(s). AbMD is used to allow the compound to rapidly explore radial space around the $\vec{r}_{pbs}$ and generate a set of structures to seed unbiased diffusive sampling. We set the AbMD mode to FLOODING with a selection frequency of 1000 and a selection constant of 0.001. The Pseudo Temperature was set to 1000K, relaxation time of 1, and resolution of 0.1 A. The collective variable is constructed by selecting two groups of atoms to define the COM distance selective variable in AMBER

#### 1.11 Hydrogen Exchange (HX or HDX)

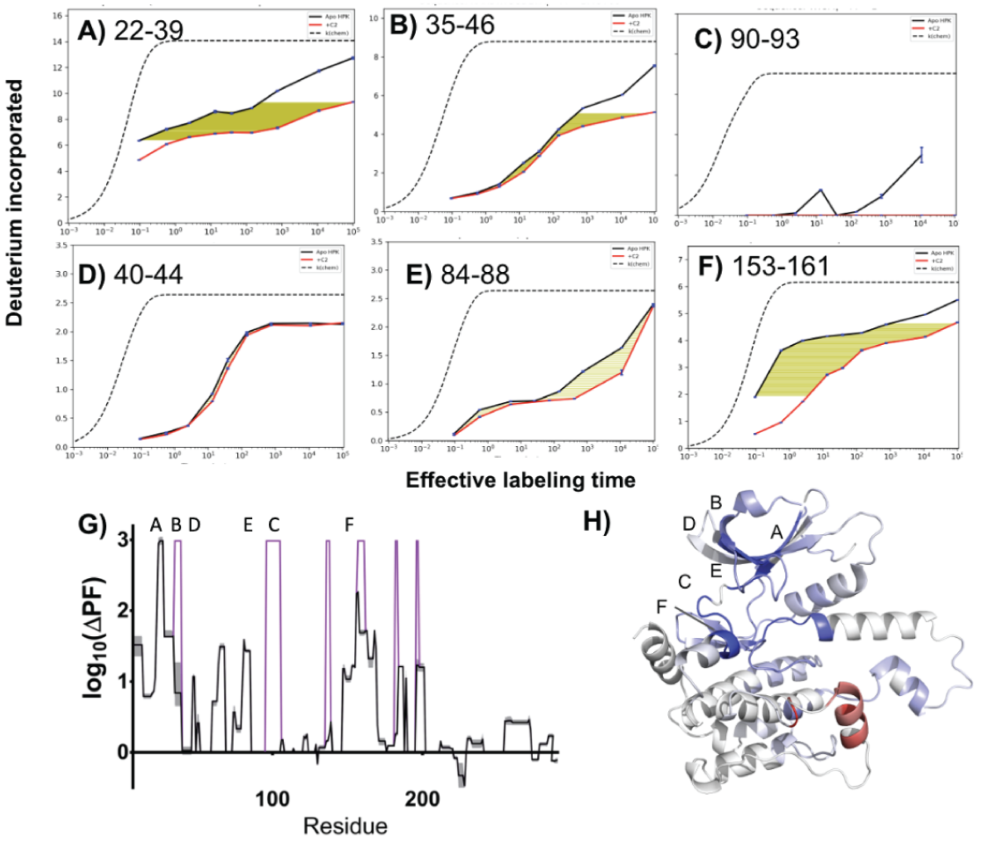


Figure 2: uptake plots with area quantified in the accompanying PF map. **A-F)** give representative uptake traces for a selection of peptides in response to binding C2 (Apo in black, bound in red) and the yellow shaded region shows the area included in computing the PF. **G)** PF plot, black trace shows the measurement and the gray shading represents uncertainty using the empirical method discussed in the text. The magenta lines show regions set to limits after manual analysis (**A** shows the largest measured PF, **B-C** are set to that value). **H)** PF changes are mapped onto the structure by $\log_{10}(\Delta PF)$. Briefly, dark blue shows the maximum log10($\Delta$PF) smaller values map linearly from blue to white, where $\log_{10}(\Delta PF)=0$. Independently, the minimum, or most negative $\log_{10}(\Delta PF)$, showing faster exchange in the bound state, is shaded dark red, also scaling linearly to white, as before.

##### 1.11.1 Sampling Philosophy

To successfully use HX-ESP, obtaining complete sequence coverage, which is a focus of most HX research [5], will not be sufficient. PFs for each residue in the protein must either be quantified or estimated, and this requires properly sampling the exchange rate of each amide. The necessary time domain can be counterintuitive and appreciated by invoking perturbation theory. To measure the perturbation, one must focus on the one being perturbed (native structure). The phenomenon of slowly exchanging protons, a focus of the first HX practitioners, was demonstrated to result from global unfolding [6], an intrinsically rare event for stable proteins. To measure perturbations on the exchange rates of these protons one needs to wait, or modulate the dominant exchange catalyst (hydroxide). Thus, for the targets interrogated in this work, two different pH values were measured iteratively, as described further in the methods enabling a large time domain to be conveniently measured in the lab. The importance of adequate sampling cannot be overstated when considering the downstream utility for HX-ESP. Had we stopped at pH 7.5, one of the largest PF perturbations in HPK1, reporting on interactions between the ligand and the ATP-pocket hinge, would not have been observed (**Extended data Figure 2c**).

###### 1.11.2 Experimental

For every compound tested, a time domain of +30 seconds to +24 hours was sampled at pH 7 and pH 9.4. Measurements at pH 9.4 were referenced to pH 7. After rescaling timepoints to a reference pH [7][8], times that are within 5 percent of the total log-scaled sampling domain are combined by averaging and this aggregated data set used for analysis. Once referenced, the effective time domain of the experiment extends from +30 seconds to roughly +100 days. PFs were then extracted using a blind empirical method that estimates a peptide-averaged PF value. This approach converges with site resolved NMR data [9], providing reliable detection of large protection factors by MS measurements. All deuterium uptake curves were manually inspected and those that diverged at early or final timepoints were flagged as potentially not properly quantified by the experimental method. These peptides were assigned the largest protection factor observed in that particular experiment. The purpose of this strategy was specifically driven towards enabling computation.

To begin the labeling experiments, 3 $\mu$L of 40 $\mu$M 30 $\mu$M for HPK-1 or PAK-1, respectively, with or without a particular compound, was diluted by the addition of 60 $\mu$L $D_2O$ buffer at either a nominal pH of 7.0 or 9.0, composed of 10 mM HEPES or ammonium acetate, and 150 mM NaCl. Given an estimated dissociation constant of 60 $\mu$M for HPK-1 [10], we expect less than 1 % of the 1.9 $\mu$M HPK-1 to exist in the dimeric form during labeling experiments. After the collection of a six-point timeseries in triplicate, logarithmically sampling a span of 30 seconds to 24 hours, samples were diluted 1:1 with a quench buffer composed of 4M GdmCl, 1M glycine, pH 2.50, and then 95 $\mu$L injected into a 0 degree (+/- 1 degree C) chamber where it is digested online with a 50:50 Fungal protease / Pepsin column (NovaBioAssays) and then loaded onto a C8 trap column (Waters, Acquity UPLC BEH C8 VanGuard Pre-column, 130Å, 1.7 µm, 2.1 mm X 5 mm) where the sample is washed for approximately 3 minutes (150 $\mu$L/min, mobile phase A) before being brought online with a C18 analytical column (Waters, ACQUITY UPLC BEH C18 Column, 130Å, 1.7 µm, 1 mm X 50 mm) and eluted by acetonitrile gradient at 35 $\mu$L/min. The eluate enters the gas phase by electro-spray (3400v, H-ESI source) and the amount of carried deuterium determined using a Thermo-Fisher Exploris 480 (120 Hz resolution, MS-only mode). Mobile phase A was water and mobile phase b, acetonitrile, each with 0.1 % formic acid and 0.04 % trifluoro acetic acid.

Using the same procedure, but with an $H_2O$ based dilution buffer and no labeling time, the peptides and associated retention times are identified by data-dependent MS/MS (top 5). This provides an initial list of retention times used by the ExMS 2.0 program [11]

For all HPK-1 measurements, the MAP4K1.D2-N293.T165E.S171E construct was used comprising the kinase domain of MAP4K1 and harboring the so-called TSEE mutations (T165E, S171E). For all PAK-1 measurements, the PAK1.S249-H545.D389N.T423E construct was used, comprising a doubly mutated (D389N, T423E) PAK-1 kinase domain. See the subsection on structural references for more information on compound disambiguation.

##### 1.12 Affinity determination

###### 1.12.1 PAK1 $K_i$

Kinase inhibition assays were described in previous work [12]

Briefly, The activity/inhibition of human recombinant PAK1 (kinase domain) or PAK2 (full length) was assessed using a Z'-LYTE™ assay (Invitrogen) by measuring the phosphorylation of a FRET peptide substrate (Ser/Thr19) labeled with Coumarin and Fluorescein. The 10 µL assay mixtures contained 50 mM HEPES (pH 7.5), 0.01 percent Brij-35, 10 mM MgCl2, 1 mM EGTA, 2 µM FRET peptide substrate, and 20 pM PAK1. Incubations were performed at 22°C in black polypropylene 384-well plates (Corning Costar).

Enzyme, FRET peptide substrate, and serially diluted test compounds were preincubated in assay buffer (7.5 µL) for 10 minutes, and the assay was initiated by adding 2.5 µL assay buffer containing 4x ATP and 160 µM PAK1. After a 60-minute incubation, the mixtures were quenched with 5 µL of Z'-LYTE™ development reagent. One hour later, emissions of Coumarin (445 nm) and Fluorescein (520 nm) were measured after excitation at 400 nm using an Envision plate reader (Perkin Elmer). The emission ratio (445 nm/520 nm) quantified substrate phosphorylation.

###### 1.12.2 HPK1 $K_i$

SLP76 phosphorylation is a proxy of HPK1 activity and inhibition of HPK1 was monitored via a SLP76 phosphorylation assay described previously [10].

Human HPK1 (full-length, T165E, S171E mutant, expressed in baculovirus, Genentech) was incubated for 30 minutes at a concentration of 0.25nM with test compounds in assay buffer (50 µM HEPES, pH 7.5/10 µM MgCl2, 2 µM TCEP, 0.01% Brij-35, 0.01% bovine serum globulins). Equal volume of substrate mix (2 mM ATP, 200 nM biotinylated SLP76) in assay buffer was added and SLP76 phosphorylation reaction carried out for 60 minutes. Phosphorylated SLP76 was detected by addition of Europium chelate-conjugated anti-phosphoSLP76(Ser376). TR-FRET signal was measured at 615 and 665 nm (Envision, Perkin Elmer) and FRET was used to calculate the % activity at each concentration of compound. $K_i$ values were calculated using the Morrison quadratic equation for tight-binding inhibitors using an ATP $K_m$ value of 20 nM.

##### 1.13 Quantitative Evaluation of Results

| Target Protein | Bound Ligand | Mean contact distance difference RMSD (Å) | | |
|---|---|---|---|---|
| | | HX-ESP | Flexible Docking | *XR Models |
| PAK1 | C1 | 0.98 ± 1.16 | 1.16 ± 1.13 | 2.07 ± 2.23 |
| *STR3448 as compute input* | C2 | 1.25 ± 1.06 | 2.55 ± 1.19 | 1.38 ± 0.94 |
| | C3 | 0.86 ± 0.61 | 5.36 ± 4.43 | 0.63 ± 0.45 |
| HPK1 | C4 | 0.58 ± 1.26 | 3.89 ± 2.34 | 0.30 ± 0.19 |
| *STR5205 as compute input* | C5 | 2.17 ± 2.54 | 8.03 ± 4.28 | 1.49 ± 1.81 |
| | C6 | 1.01 ± 1.06 | 9.27 ± 4.54 | 1.13 ± 1.58 |

Table 1: Mean contact distance difference RMSD

This table reports the percentage of models that contain the particular interatomic contact observed in the X-ray structure
**Hbonds are bold, Hinge H-bonds are bold red**
Cutoffs: 4.5 angstrom (non polar contacts); 3.5 angstrom (H-bond)

**HPK - C3**

| Contacts | HX-ESP | HADDOCK | X-ray+MD |
|---|---|---|---|
| 23 | 80 | 0 | 100 |
| 24 | 80 | 0 | 100 |
| 28 | 60 | 0 | 67 |
| 31 | 60 | 0 | 67 |
| 44 | 100 | 0 | 100 |
| 91 | 100 | 0 | 67 |
| **92** | **100** | **0** | **100** |
| 93 | 60 | 100 | 100 |
| **94** | **100** | **100** | **100** |
| 95 | 40 | 0 | 0 |
| 96 | 40 | 0 | 100 |
| 97 | 100 | 100 | 100 |
| 144 | 80 | 0 | 100 |

**HPK - C4**

| Contacts | HX-ESP | HADDOCK | X-ray+MD |
|---|---|---|---|
| 23 | 100 | 0 | 100 |
| 24 | 100 | 0 | 100 |
| 28 | 40 | 0 | 100 |
| 31 | 100 | 0 | 100 |
| 44 | 100 | 0 | 100 |
| 75 | 100 | 0 | 100 |
| 91 | 100 | 0 | 100 |
| **92** | **100** | **0** | **100** |
| 93 | 100 | 0 | 100 |
| **94** | **100** | **0** | **100** |
| 95 | 100 | 0 | 100 |
| 97 | 100 | 0 | 100 |
| **101** | **100** | **0** | **100** |
| 144 | 100 | 100 | 100 |
| 154 | 100 | 0 | 100 |
| **155** | **100** | **0** | **100** |

**HPK - C5**

| Contacts | HX-ESP | HADDOCK | X-ray+MD |
|---|---|---|---|
| **21** | **0** | **0** | **33** |
| 23 | 100 | 0 | 67 |
| 28 | 0 | 0 | 0 |
| 31 | 0 | 0 | 67 |
| 44 | 100 | 0 | 67 |
| **46** | **0** | **0** | **100** |
| 75 | 40 | 0 | 33 |
| **91** | **60** | **0** | **100** |
| **92** | **100** | **0** | **100** |
| 93 | 100 | 0 | 100 |
| **94** | **100** | **0** | **100** |
| **95** | **40** | **0** | **33** |
| 97 | 100 | 0 | 67 |
| 101 | 20 | 0 | 33 |
| 142 | 0 | 0 | 33 |
| 144 | 100 | 0 | 100 |
| 156 | 0 | 0 | 0 |

**HPK - C6**

| Contacts | HX-ESP | HADDOCK | X-ray+MD |
|---|---|---|---|
| 23 | 20 | 0 | 100 |
| 31 | 60 | 0 | 0 |
| 44 | 100 | 0 | 100 |
| 46 | 100 | 0 | 100 |
| 75 | 80 | 0 | 100 |
| 91 | 60 | 0 | 67 |
| **92** | **100** | **0** | **100** |
| 93 | 100 | 25 | 100 |
| **94** | **100** | **0** | **100** |
| **101** | **40** | **0** | **67** |
| 144 | 100 | 0 | 0 |

**PAK1 - C1**

| Contacts | HX-ESP | HADDOCK | X-ray+MD |
|---|---|---|---|
| 297 | 100 | 100 | 100 |
| **299** | **100** | **0** | **100** |
| 315 | 40 | 75 | 33 |
| 316 | 60 | 0 | 33 |
| 328 | 0 | 0 | 33 |
| 342 | 100 | 0 | 100 |
| 344 | 100 | 50 | 100 |
| 345 | 100 | 100 | 100 |
| 346 | 80 | 100 | 67 |
| **347** | **100** | **100** | **100** |
| 350 | 100 | 100 | 67 |
| 393 | 100 | 100 | 67 |
| 394 | 0 | 0 | 0 |
| 396 | 80 | 0 | 0 |
| 406 | 80 | 100 | 0 |
| **407** | **80** | **75** | **0** |

**PAK1 - C2**

| Contacts | HX-ESP | HADDOCK | X-ray+MD |
|---|---|---|---|
| 284 | 0 | 0 | 0 |
| 297 | 80 | 0 | 100 |
| 299 | 0 | 0 | 33 |
| 301 | 40 | 0 | 0 |
| 315 | 0 | 50 | 0 |
| 344 | 100 | 0 | 67 |
| 345 | 100 | 0 | 100 |
| 346 | 100 | 0 | 67 |
| **347** | **40** | **0** | **100** |
| 396 | 80 | 0 | 33 |
| 406 | 80 | 75 | 33 |

Table 2: Percentage of models that contain correct contacts to the target protein

**HPK - C1**

| Contacts | HX-ESP | HADDOCK | X-ray+MD |
|---|---|---|---|
| 23 | 3.8 | 7.8 | 3.8 |
| 24 | 3.6 | 6.8 | 3.6 |
| 28 | 4.5 | 8.4 | 4.4 |
| 31 | 4.6 | 11.4 | 4.3 |
| 44 | 3.5 | 10.1 | 3.4 |
| 91 | 3.8 | 19.8 | 4.3 |
| **92** | **3.4** | **10.8** | **3.5** |
| 93 | 4.3 | 3.8 | 4.2 |
| **94** | **3.0** | **2.8** | **3.1** |
| 95 | 4.7 | 8.0 | 4.9 |
| 96 | 5.3 | 6.6 | 3.8 |
| 97 | 3.7 | 3.9 | 3.7 |
| 144 | 3.9 | 12.6 | 3.8 |

**HPK - C2**

| Contacts | HX-ESP | HADDOCK | X-ray+MD |
|---|---|---|---|
| 23 | 3.9 | 9.6 | 4.0 |
| 24 | 3.9 | 8.9 | 3.9 |
| 28 | 6.4 | 11.6 | 3.8 |
| 31 | 3.9 | 12.7 | 3.6 |
| 44 | 3.6 | 5.5 | 3.7 |
| 75 | 3.8 | 4.9 | 3.9 |
| 91 | 3.6 | 7.0 | 3.7 |
| **92** | **2.9** | **5.6** | **3.0** |
| 93 | 3.7 | 7.3 | 3.7 |
| **94** | **3.1** | **6.2** | **3.2** |
| 95 | 4.0 | 8.0 | 3.5 |
| 97 | 3.8 | 5.6 | 3.7 |
| **101** | **3.5** | **7.4** | **3.1** |
| 144 | 3.6 | 4.2 | 3.7 |
| 154 | 3.8 | 5.5 | 3.9 |
| **155** | **3.0** | **6.4** | **3.3** |

**HPK - C3**

| Contacts | HX-ESP | HADDOCK | X-ray+MD |
|---|---|---|---|
| **21** | **6.0** | **15.8** | **4.7** |
| 23 | 4.1 | 5.5 | 4.3 |
| 28 | 12.3 | 12.7 | 8.2 |
| 31 | 7.1 | 9.8 | 4.0 |
| 44 | 3.7 | 8.2 | 4.1 |
| **46** | **6.4** | **16.2** | **3.1** |
| 75 | 4.7 | 12.6 | 4.8 |
| **91** | **3.5** | **16.4** | **3.4** |
| **92** | **3.1** | **8.3** | **2.8** |
| 93 | 3.9 | 9.4 | 3.8 |
| **94** | **2.8** | **6.5** | **3.0** |
| **95** | **3.6** | **13.1** | **3.7** |
| 97 | 3.8 | 6.9 | 3.8 |
| 101 | 5.9 | 11.4 | 6.0 |
| 142 | 7.8 | 14.6 | 4.9 |
| 144 | 3.7 | 6.4 | 3.8 |
| 156 | 8.9 | 19.6 | 9.9 |

**HPK - C4**

| Contacts | HX-ESP | HADDOCK | X-ray+MD |
|---|---|---|---|
| 23 | 5.6 | 11.4 | 3.7 |
| 31 | 5.3 | 17.0 | 7.6 |
| 44 | 3.8 | 11.1 | 3.6 |
| 46 | 3.5 | 19.6 | 3.9 |
| 75 | 4.3 | 14.2 | 4.2 |
| 91 | 4.1 | 12.8 | 4.7 |
| **92** | **2.8** | **10.0** | **2.8** |
| 93 | 3.7 | 4.8 | 3.7 |
| **94** | **3.0** | **7.7** | **3.0** |
| **101** | **4.9** | **18.2** | **3.1** |
| 144 | 3.7 | 11.8 | 6.9 |

**PAK1 - C1**

| Contacts | HX-ESP | HADDOCK | X-ray+MD |
|---|---|---|---|
| 297 | 4.0 | 4.4 | 4.0 |
| **299** | **2.8** | **7.5** | **2.8** |
| 315 | 5.4 | 4.3 | 4.6 |
| 316 | 4.5 | 6.4 | 6.0 |
| 328 | 5.0 | 5.5 | 5.8 |
| 342 | 3.5 | 4.8 | 3.8 |
| 344 | 3.8 | 4.5 | 3.9 |
| 345 | 3.7 | 4.1 | 3.7 |
| 346 | 3.8 | 3.5 | 4.1 |
| **347** | **2.8** | **2.8** | **2.8** |
| 350 | 3.8 | 3.5 | 5.2 |
| 393 | 3.8 | 3.3 | 4.9 |
| 394 | 8.0 | 5.6 | 9.2 |
| 396 | 4.2 | 4.8 | 5.5 |
| 406 | 4.1 | 3.3 | 6.4 |
| **407** | **3.2** | **3.5** | **10.7** |

**PAK1 - C2**

| Contacts | HX-ESP | HADDOCK | X-ray+MD |
|---|---|---|---|
| 284 | 4.9 | 6.9 | 5.2 |
| 297 | 4.2 | 9.1 | 4.0 |
| 299 | 5.1 | 5.7 | 4.6 |
| 301 | 5.1 | 6.0 | 5.1 |
| 315 | 6.5 | 4.4 | 6.2 |
| 344 | 3.8 | 6.5 | 4.4 |
| 345 | 3.7 | 5.4 | 3.5 |
| 346 | 3.6 | 5.4 | 4.4 |
| **347** | **3.9** | **5.7** | **2.8** |
| 396 | 4.3 | 5.8 | 5.0 |
| 406 | 3.7 | 4.5 | 5.4 |

Table 3: Mean distance across all models for each contact to the target protein

##### 1.13.1 Contact distance Difference RMSD

In order to draw a standardized comparison between various structural poses, we employed the strategy of formulating a contact distance difference matrix. This matrix is composed of atomic contact pairs, each representing the interaction between the ligand and the protein. In the course of defining each contact pair, we identify symmetric atoms within individual molecules and regard them as equal entities. One significant advantage of this method is its ability to maintain invariance, irrespective of fluctuations in a carbon alpha Root Mean Square Deviation (RMSD) alignment. This characteristic facilitates a precise identification of binding modes within inherently dynamic systems, and this is accomplished without penalizing positional or locally equivalent variability. Although the process of identifying and segregating symmetric atoms can be time-consuming, it is made feasible thanks to the remapping algorithm, see **Extended data Table 3** , which autonomously performs this task.

In evaluating contact difference RMSD, we utilize a reference structure with $M$ contacts between protein atoms $\vec{A}$ and ligand atoms $\vec{B}$. The elements of $\vec{A}$ and $\vec{B}$ are atom positions and are paired by index. We then construct a contact difference vector $\vec{R}$:

$$R_i = \left|\vec{A}_i - \vec{B}_i\right|$$

In comparing $\vec{R}$ across different structures, we construct the reference structure's contact distances $\vec{R}_{\mathrm{ref}}$ and the target structure's contact distances $\vec{R}_{\mathrm{tar}}$. Given that the $N$ input target models share the same connectivity, we construct a contact difference matrix $\boldsymbol{D}$:

$$\boldsymbol{D} = \left[\vec{R}_{\mathrm{tar}0} - \vec{R}_{\mathrm{ref}}, \ldots, \vec{R}_{\mathrm{tar}N} - \vec{R}_{\mathrm{ref}}\right]$$

We then construct the contact difference RMSD vector $\vec{\sigma}$:

$$\vec{\sigma}_i = \sqrt{\left\langle \boldsymbol{D}_i^2 \right\rangle - \left\langle \boldsymbol{D}_i \right\rangle^2}$$

With $\vec{\sigma}$, we relate the target models to a mean RMSD value and an uncertainty as

$$\overline{\mathrm{RMSD}} = \langle \vec{\sigma} \rangle$$

$$\sigma_{\overline{\mathrm{RMSD}}} = \sqrt{\langle \vec{\sigma}^2 \rangle - \langle \vec{\sigma} \rangle^2}$$

##### 1.13.2 Alignments for pocket RMSD and ligand RMSD

For atomic RMSD calculations (Main Text **Table 1**) and figures that represent structural overlays ( **main text figures 3-5**), first all structures are aligned with the input structure apo HPK-1 or apo PAK-1 used for docking or HX-ESP computations. Using the word "MODEL" to represent a predicted binding pocket result (HX-ESP or HADDOCK), and for MD relaxed X-ray structures, "TARGET" to represent the bound X-ray structure, and "INPUT" to represent the respective unbound X-ray structure, the following pymol commands were used to produce overlays and subsequently calculate the pocket and ligand RMSD. The same "'super"' command was used to produce all alignments shown in the manuscript.

*Pocket RMSD - Target to input*:
```
super TARGET and byres resn LIG around 8 and (name n or name ca or name c or name o), INPUT
```

*Pocket RMSD - Model to input or target*:
```
super MODEL and byres resn LIG around 8 and (name n or name ca or name c or name o), [INPUT
or TARGET]
```

*Ligand RMSD - Model to target*:
```
rms_cur MODEL and resn LIG, TARGET and resn LIG, matchmaker=-1
```

#### 1.14 Crystal Structures

##### 1.14.1 PAK1 Structures

EDITOR NOTE: Prior to submission, the identifiers will be removed.

Apo Structure - STR3448 - G02854177 - PDBID:TBD
C1 - STR3800 - G02857900 - PDBID:TBD
C2 - STR3432 - G02854702 - PDBID:TBD

The PAK1 crystal structures (Extended data Table 4) of these inhibitors at resolution 1.8-2.3Å showed all of the compounds bound at the structurally well-studied ATP binding site[13][14]. The amino acid positions observed in each inhibitor complex remained largely unchanged (Rmsd range 0.27-0.58Å for all structures in this study, main text Table 1). The electron density for the three compounds with the highest affinity fully covered all inhibitor atoms (Extended data Figure 3a). In each inhibitor complex, multiple hydrogen bonds were formed between the ligand and the hinge region of the kinase (amino acid backbone positions 345 and 347). Inhibitors G02854177 and C1 formed additional hydrogen bonds with the Lys299 sidechain, and G02854177 formed an ion pair with the sidechain of Asp407 from its tertiary amine tail (Extended data Figure 3b). Each inhibitor also engaged in van der Waals interactions with multiple protein residues in the binding site. The structure of G02854177 was selected as the template for docking studies, since it was the highest resolution complex.

The unliganded input structure was derived from STR3448. The ligand was manually removed from this structure, the structure was protonated at pH 7.5 using proPKA. A series of two simulations was run to eliminate the structural memory of the deleted compound, first AMD, and then cMD. The most abundant structure was propagated from AMD to CMD using the same k-means algorithm used throughout the work. This created the input structure file. ACE and NME caps were placed on N- and C- termini due to their proximity to the binding pocket.

##### 1.14.2 HPK1 Structures

EDITOR NOTE: Prior to submission, the identifiers will be removed.

Apo Structure - STR5205 - G03089260 - PDBID: TBD
C3 - STR3656 - G02931858 - PDBID:TBD
C4 - STR6214 - G03427872 - PDBID:TBD
C5 - STR6179 - G03407781 - PDBID:TBD
C6 - STR6102 - G03418184 - PDBID:TBD

The structures of HPK1 (Extended data Table 5) all were determined of the previously described [10] T165E/S171E mutant, emulating activated protein. Unlike the PAK1 structures, the HPK1 structures showed marked conformational differences in their P-loops (aka G-loops) and activation helices and the relative orientations of the N- and C-lobes of the kinase domain, derived from a combination of ligand- and (potentially) lattice-driven influences. The compound originally bound to the STR5205 complex structure, used as the input in these docking studies and the complexes with C3 and C4 all exhibited classical Type I hinge binding, with the DFG motif in the "in" conformation. The complex with C5 bound as Type 1.5 mode, with the DFG motif "out" but the compound remaining entirely on the front side of it, along the hinge, and not entering the back pocket. The final complex of HPK1 with C6, showed the largest conformational changes, relative to the other structures, with the DFG motif significantly extended and the activation helix nearly perpendicular to that of the other structures. This complex had multiple binding sites occupied by inhibitors, with both chains having a Type 1.5 binding mode, with the compound at the hinge. Additionally, chain A had an inhibitor molecule bound in a pocket under the DFG motif and adjacent to helix 3, a variation on the Type III binding mode. It is unclear, if this site is relevant to the pharmacology in the cell or is a product of the lattice. It was not seen in the prior structure of a similar compound (PDBID: 8PAS[15]). Additionally, we found no evidence of binding at this site by HX experiments and therefore considered only the Type 1.5 binding mode in benchmarking exercises. Two other inhibitor sites were also present, one at the lattice contact of the two activation helices and the other in the crevice of above the N-lobe. Both of these binding modes would only occur because of the crystal lattice. All of the ligands showed strong electron density (Extended data Figure (3b)) covering the compounds, except for C6, that had a disordered acyclic tail and pendant morpholine moiety. The terminal nitrile of C5 also had weak density.

The unliganded input structure was derived from STR5205. To prepare this file for molecular dynamics or docking, the compound was manually deleted and the file was protonated using proPKA. A series of two simulations were run to eliminate structural memory of the deleted compound, first AMD, and then CMD. The most abundant structure was propagated from AMD to CMD using the same k-means algorithm used throughout the work. This created the input structure file.

##### 1.15 Protein expression, purification, crystallization, and structure determination

The PAK1 and HPK1 kinases were expressed, purified, and crystallized as previously described [16][10]. Crystallographic data were collected at beamlines indicated in Extended data Tables 4 and 5. Data were integrated scaled with HKL2000 [17] or XDS [18], and the structures were determined by molecular replacement with PHASER [19]. The models were built in COOT [20] and subsequently refined with PHENIX [21] to final statistics presented in Extended data Tables 4 and 5. The simulated annealing omit difference electron density maps (Extended data Figure 3) demonstrate the fidelity of the final models with the electron density.

| | Apo - G02854177 STR3448 | C1 - G02854702 STR3432 | C2 - G02857900 STR3800 |
|---|---|---|---|
| PDB code | AAA | BBB | CCC |
| X-ray source | APS 21-ID-G | APS 21-ID-G | CRO |
| Wavelength (Å) | 0.97856 | 0.97856 | - |
| Detector | Rayonix M300 | Rayonix M300 | - |
| Resolution range (Å) | 50.0 – 1.78 | 50.0 – 1.85 | 50.0 – 2.15 |
| Highest res. bin (Å) | 1.84 – 1.78 | 1.92 – 1.85 | 2.19 – 2.15 |
| Space group | P21 | P21 | P21 |
| *Multiplicity | 3.8, 3.8 | 3.8, 3.6 | 3.1, 2.6 |
| *Complete (%) | 99.1, 98.3 | 98.3, 97.5 | 97.8, 97.6 |
| *Mean $I/s_I$ | 24.5, 2.1 | 21.5, 2.1 | 12.7, 1.9 |
| Wilson B ($Å^2$) | 28.1 | 29.2 | 48.4 |
| CC $\frac{1}{2}$ (highest bin) | N/A | N/A | 0.852 |
| $R_{merge}$ (%) | 4.9, 69.3 | 5.7, 66.7 | 7.3, 50.8 |
| **Refinement** | | | |
| # reflections ($R_{free}$ set) | 59,824 (3,022) | 52,866 (2,694) | 33,297 (1,662) |
| Resolution range (Å) | 30.6 – 1.78 | 34.1 – 1.85 | 31.9 – 2.15 |
| $R_{work}$, $R_{free}$ (%) | 17.2, 20.3 | 17.6, 21.6 | 22.5, 26.5 |
| # non-H atoms | 4,472 | 4,486 | 4,239 |
| # solvent molecules | 489 | 388 | 41 |
| Rmsd bond lengths (Å) | 0.013 | 0.006 | 0.006 |
| Rmsd bond angles (rad) | 1.33 | 0.916 | 0.717 |
| Ramachandran fav. (%) | 97.7 | 96.8 | 96.1 |
| Ramachand. outlier (%) | 0 | 0.2 | 0.4 |
| Ave B-factor ($Å^2$) | 43.8 | 41.7 | 62.3 |
| Molprobity clash score | 3.6 | 3.3 | 8.7 |

*values for the full and highest resolution ranges are comma separated

Table 4: Crystallographic data for PAK1 structures

| | Apo - G03089260 STR5205 | C3 - G02931858 STR3656 | C4 - G03427872 STR6214 | C5 - G03407781 STR6179 | C6 - G03418184 STR6102 |
|---|---|---|---|---|---|
| PDB code | XXX | YYY | ZZZ | QQQ | RRR |
| X-ray source | APS 21-ID-F | APS 21-ID-F | APS 21-ID-D | SSRF BL18U1 | ALS 8.3.1 |
| Wavelength (Å) | 0.97872 | 0.97872 | 0.97918 | 0.97853 | 1.11583 |
| Detector | Rayonix M-225 | Rayonix M-225 | Eiger 9M | Pilatus3 6M | Pilatus3 6M |
| Resolution range (Å) | 52.47 – 1.80 | 42.47 – 1.50 | 44.63 – 1.45 | 44.50 – 2.30 | 47.94 – 2.13 |
| Highest res. bin (Å) | 1.89 – 1.80 | 1.59 – 1.50 | 1.47 – 1.45 | 2.38 – 2.30 | 2.19 – 2.13 |
| Space group | P1 | P1 | C2 | P21 | P21 |
| *Multiplicity | 2.0, 2.0 | 2.2, 2.2 | 7.2, 6.9 | 3.3, 3.4 | 6.7, 6.9 |
| *Complete (%) | 97.1, 95.9 | 93.9, 94.1 | 94.9, 85.3 | 99.6, 99.3 | 98.8, 99.2 |
| *Mean $I/s_I$ | 9.8, 2.2 | 11.7, 3.2 | 13.4, 3.6 | 8.9, 2.1 | 14.1, 2.7 |
| Wilson B ($Å^2$) | 19.9 | 12.4 | 14.7 | 49.0 | 34.6 |
| CC $\frac{1}{2}$ (highest bin) | 0.697 | 0.883 | 0.881 | 0.791 | 0.817 |
| $R_{merge}$ (%) | 4.8, 37.4 | 3.6, 24.6 | 7.4, 61.1 | 13.1, 63.9 | 8.8, 81.1 |
| **Refinement** | | | | | |
| # reflections ($R_{free}$ set) | 54,142 (2,675) | 89,881 (4,474) | 193,687 (9,367) | 24,210 (1,234) | 34,675 (1,699) |
| Resolution range (Å) | 52.5 – 1.80 | 27.5 – 1.50 | 33.8 – 1.45 | 35.0 – 2.30 | 47.9 – 2.13 |
| $R_{work}$, $R_{free}$ (%) | 15.9, 20.0 | 15.6, 18.0 | 16.5, 18.4 | 22.5, 27.6 | 17.8, 22.7 |
| # non-H atoms | 4,606 | 4,537 | 4,602 | 4,709 | 4,561 |
| # solvent molecules | 561 | 766 | 574 | 110 | 232 |
| Rmsd bond lengths (Å) | 0.005 | 0.005 | 0.007 | 0.003 | 0.005 |
| Rmsd bond angles (rad) | 1.299 | 0.842 | 0.951 | 0.925 | 0.774 |
| Ramachandran fav. (%) | 97.8 | 98.1 | 97.6 | 96.2 | 97.7 |
| Ramachand. outlier (%) | 1.0 | 0 | 0 | 0.2 | 0.2 |
| Ave B-factor ($Å^2$) | 27.8 | 19.3 | 21.1 | 51.7 | 41.4 |
| Molprobity clash score | 3.3 | 3.0 | 3.0 | 3.7 | 1.5 |

*values for the full and highest resolution ranges are comma separated

Table 5: Crystallographic data for HPK1 structures

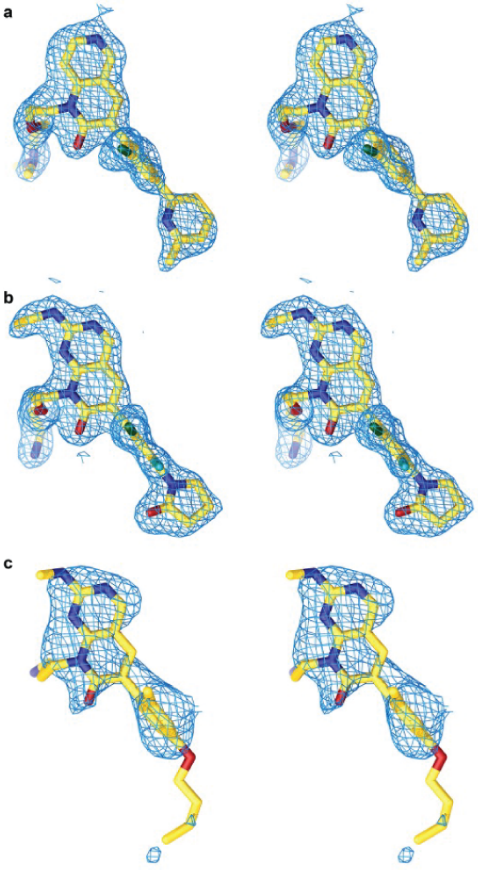


(a) PAK1 electron density maps. Divergent eye stereo images of the simulated annealing composite omit difference electron density maps, (2m—Fo— - D—Fc—) exp(iαc), of the PAK1 inhibitor complexes G02854177 (**a**), C2 (**b**), C1 (**c**), contoured at 1s.

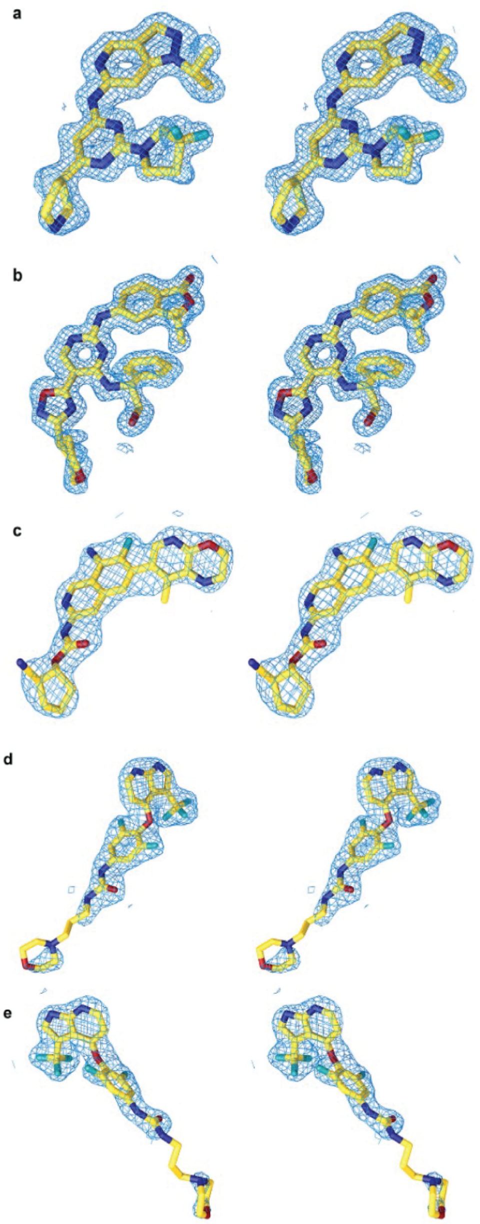


(b) HPK1 electron density maps. Divergent eye stereo images of the simulated annealing composite omit difference electron density maps, (2m—Fo— - D—Fc—) exp(iαc), of the HPK1 inhibitor complexes C3 (**a**), C4 (**b**), C5 (**c**), C6 hinge site (**d**) and Type III-like (**e**) site, contoured at 1s.

Figure 3

#### 1.16 PBS composition

Note, the PBS numbers listed below refer to linear numbered structure files made available in our GIT repository. The offsets need to correct these numbers to the canonical numbering scheme found in the manuscript are as follows:

All PBS residues refer to positions in the apo structure. In one case for HPK-1 and PAK-1, the target X-ray structures have a different sequence than the apo structures used for all simulations. The offsets below only apply to the provided apo structures.

MD PAK1 + 248 = Canonical PAK1

MD HPK1 + 2 = Canonical HPK1

If the PBS needed to be modified for running the WSP algorithm (**Extended data Figure 1**) the modified PBS has been listed in addition to the HX-ESP/HADDOCK PBS. This is a notable weakness of the WSP algorithm, and does rely on subjective interpretation from the operator of HX-ESP. It can be substituted, however, by using HADDOCK, or any other docking program to generate starting structures.

##### 1.16.1 PAK1:C1

PBS: 39, 84, 85, 100, 101, 146, 147, 148, 149, 160, 161
WSP PBS: 28,29,39,40,77,78,79,80,81,82,83,84,85,86,87,88,100,131,138,139,140,141,142, 30,31,32,33,34,35

##### 1.16.2 PAK1:C2

PBS: 28, 29, 31, 33, 34, 36, 37, 38, 39, 40, 51, 52, 53, 54, 55, 56, 65, 66, 67, 68, 69, 70, 71, 72, 73, 81, 82, 83, 84, 100, 101, 144, 145, 146, 147, 148, 149, 160, 161, 163
The same PBS was used for WSP.

##### 1.16.3 HPK1:C3

PBS: 19, 20, 21, 25, 26, 28, 29, 30, 43, 44, 70, 71, 92, 94, 96, 97, 98
The same PBS was used for WSP.

##### 1.16.4 HPK1:C4

PBS:20, 21, 31, 32, 33, 34, 35, 92, 94, 96, 97, 98, 99, 100, 101, 153
The same PBS was used for WSP.

##### 1.16.5 HPK1:C5

PBS: 20, 21, 31, 32, 33, 65, 66, 67, 68, 71, 94, 96, 97, 98, 99, 100, 152, 153, 154
The same PBS was used for WSP.

##### 1.16.6 HPK1:C6

PBS: 20, 21, 31, 32, 33, 72, 73, 74, 75, 92, 94, 96, 97, 99, 100
The same PBS was used for WSP.

#### 1.17 X-ray relaxed Models

To produce models of the target structures that show whether contacts observed in the target structures are expected to be affected by refolding of domain swapped elements, X-ray structures were subjected to three replicates of conventional MD at 375K (HPK-1) and 350K (PAK-1) for 1 microsecond. The K-means selection algorithm, combined with the PBS, was used to select a single representative model from each trajectory. Each model was then simulated for 100 nanoseconds at 298K before selecting single models by the same K-means algorithm. This produced three X-ray relaxed models for each target structure to be compared with HX-ESP predictions.

##### 1.18 Identification of contacts in target structures

Contacting pairs of atoms in target X-ray structures were identified using the OpenEye OEChem TK 2022.2.1 (OpenEye, Cadence Molecular Sciences, Santa Fe, NM. http://www.eyesopen.com). Specifically, the OEInteractionHintContainer is used to perceive contacts between atoms in the ligand and residues in the protein. Once a contacting residue has been identified, the distances of the contact pairs identified by selecting the pair with the smallest distance. Contacting pairs of atoms are listed in each subsection, below.

The contacts follow the format:
:LIGAND@ATOM :PROTEIN@ATOM

Symmetry pairs identified by the remapping algorithm outlined in section 2 are separated by a comma and they can occur on the protein (eg. Phenylalanine CE1 and CE2) or on the ligand (eg. F1 and F2 on HPK1:C1). These symmetry pairs must be identified as each atom has a unique name. If a ring flips, for example, then our analysis would need to modify the contacting pair of atoms. In cases where symmetry pairs are identified, the reported distance in calculations takes the shorter of the two distances.

The following lists use residue numbers and atom names in the files that are hosted on our git repo. Please see those structures for further inquiry.

###### 1.18.1 PAK1:C1

:LIG@C4 :297@CB
:LIG@O3 :299@NZ
:LIG@C22 :315@OE2
:LIG@C21 :316@CG1
:LIG@C4 :328@CG1
:LIG@F1 :342@CG1
:LIG@C18 :344@CG
:LIG@C4 :345@O
:LIG@N1 :346@CE1,:346@CE2
:LIG@N1 :347@O
:LIG@C1 :350@CA
:LIG@N2 :396@CD1
:LIG@C15 :406@OG1
:LIG@C13 :407@OD2

###### 1.18.2 PAK1:C2

:LIG@C8 :284@CG1
:LIG@C8 :297@CB
:LIG@C4 :299@CD
:LIG@C1 :301@CE
:LIG@C1 :315@OE2
:LIG@C6 :344@CG
:LIG@C16 :345@O
:LIG@C18 :346@CE1,:346@CE2
:LIG@N2 :347@O
:LIG@C16 :396@CD1
:LIG@C10 :406@OG1

##### 1.18.3 HPK1:C3

:LIG@F1,:LIG@F2 :23@C
:LIG@F1,:LIG@F2 :24@N
:LIG@F1,:LIG@F2 :28@OH
:LIG@F1,:LIG@F2 :28@OH
:LIG@F1,:LIG@F2 :31@CB
:LIG@C6 :44@CB
:LIG@N2 :91@CE
:LIG@C6 :92@O
:LIG@N4 :93@CE1,:93@CE2
:LIG@N4 :94@O
:LIG@C18 :95@O
:LIG@C21 :96@O
:LIG@C11 :97@CA
:LIG@C5 :144@CD1

##### 1.18.4 HPK1:C4

:LIG@C19,:LIG@C21 :23@C
:LIG@C19,:LIG@C21 :24@N
:LIG@C20 :28@OH
:LIG@C19,:LIG@C21 :31@CB
:LIG@N1 :44@CB
:LIG@C6 :75@CG2
:LIG@C6 :91@SD
:LIG@N1 :92@O
:LIG@C12 :93@CD1,:93@CD2
:LIG@N2 :94@N
:LIG@N6 :95@O
:LIG@N5 :97@CA
:LIG@O3 :101@OD2
:LIG@C8 :144@CD1
:LIG@O2 :154@CB
:LIG@O2 :155@N

##### 1.18.5 HPK1:C5

:LIG@N5 :21@NE2
:LIG@C11 :23@CD1
:LIG@N1 :28@O
:LIG@C5 :31@CG2
:LIG@N3 :44@CB
:LIG@N1 :46@NZ
:LIG@N3 :75@CG1
:LIG@F1 :91@CE
:LIG@N3 :92@O
:LIG@N4 :93@CE1,:93@CE2
:LIG@N4 :94@O
:LIG@C21 :95@O
:LIG@O2 :97@CA
:LIG@C18 :101@CG
:LIG@N6 :142@OD1

:LIG@C13 :144@CD1
:LIG@N1 :156@CE1,:156@CE2

###### 1.18.6 HPK1:C6

:LIG@C8 :23@CD1
:LIG@F1 :31@CB
:LIG@C10 :44@CB
:LIG@F2,:LIG@F3,:LIG@F4,:LIG@F3,:LIG@F4 :46@CE
:LIG@C11 :75@CG1
:LIG@F2,:LIG@F3,:LIG@F4 :91@CE
:LIG@N2 :92@O
:LIG@N1 :93@CD1,:93@CD2
:LIG@N1 :94@N
:LIG@N3 :101@OD2
:LIG@F5 :144@CD2

##### 1.19 Chemical synthesis of compounds

#### 1.19.1 C1

Chemical name: 8-(2-(2-aminoethoxy)ethyl)-6-(2-chloro-3-fluoro-4-(2-oxopyrrolidin-1-yl)phenyl)-2-(ethylamino)pyrido[2,3-d]pyrimidin-7(8H)-one
Exact synthetic procedure described in reference [22].

#### 1.19.2 C2

Chemical name: 8-(3-aminopropyl)-6-(4-butoxy-2-methylphenyl)-2-(methylamino)pyrido[2,3-d]pyrimidin-7(8H)-one
Exact synthetic procedure described in reference [22]

#### 1.19.3 C3

Chemical name: N-(2-(3,3-difluoropyrrolidin-1-yl)-6-(pyrrolidin-3-yl)pyrimidin-4-yl)-1-isopropyl-1H-pyrazolo[4,3-c]pyridin-6-amine
Synthetic procedures described in reference [10]

#### 1.19.4 C4

Chemical name: 5-[(4-[(1S)-2-hydroxy-1-phenylethyl]amino-5-[3-(oxan-4-yl)-1,2,4-oxadiazol-5-yl]pyrimidin-2-yl)amino]-3,3-dimethyl-1,3-dihydro-2-benzofuran-1-one
Exact synthetic protocol with characterization has been disclosed in patent: WO2019090198 A1

#### 1.19.5 C5

Chemical name: (1R,2S)-2-cyanocyclopentyl (8-amino-7-fluoro-6-(8-methyl-2,3-dihydro-1H-pyrido[2,3-b][1,4]oxazin-7-yl)isoquinolin-3-yl)carbamate
The synthesis of C5 was performed according to the general procedure described previously [23].

Step 1:(±)-tert-butyl 7-[8-(tert-butoxycarbonylamino)-3-[[2-cyanocyclopentoxy]carbonylamino]-7-fluoro-6-isoquinolyl]-8-methyl-2,3-dihydropyrido[2,3-b][1,4]oxazine-1-carboxylate

A solution of tert-butyl 7-[3-amino-8-(tert-butoxycarbonylamino)-7-fluoro-6-isoquinolyl]-8-methyl-2,3-dihydropyrido[2,3-b][1,4]oxazine-1-carboxylate (300 mg, 0.57 mmol) and (±)-(trans)-2-hydroxycyclopentanecarbonitrile (300 mg, 2.7 mmol) in dichloromethane (60 mL) was added DIEA (500 mg, 3.88mmol) at room temperature. Triphosgene (200 mg, 0.67 mmol) was then added and stirred at 0 °C for 2 h. The residue was purified by flash chromatography on silica gel eluting with petroleum ether/ethyl acetate (1/1) to afford (±)-tert-butyl 7-[8-(tert-butoxycarbonylamino)-3-[[2-cyanocyclopentoxy]carbonylamino]-7-fluoro-6-isoquinolyl]-8-methyl-2,3-dihydropyrido[2,3-b][1,4]oxazine-1-carboxylate (220 mg, 0.332 mmol, 58.2% yield) as a yellow solid. LCMS (ESI) $[M+H]^+$ = 663.

Step 2: [(1R,2S)-2-cyanocyclopentyl] N-[8-amino-7-fluoro-6-(8-methyl-2,3-dihydro-1H-pyrido[2,3-b][1,4]oxazin-7-yl)-3-isoquinolyl]carbamate (C5) and [(1S,2R)-2-cyanocyclopentyl] N-[8-amino-7-fluoro-6-(8-methyl-2,3-dihydro-1H-pyrido[2,3-b][1,4]oxazin-7-yl)-3-isoquinolyl]carbamate (enantiomer of C5)

A solution of (±)-tert-butyl 7-[8-(tert-butoxycarbonylamino)-3-[[2-cyanocyclopentoxy]carbonylamino]-7-fluoro-6-isoquinolyl]-8-methyl-2,3-dihydropyrido[2,3-b][1,4]oxazine-1-carboxylate (220 mg, 0.33 mmol) in dichloromethane (10 mL) was added 2,2,2-trifluoroacetic acid (2 mL) at room temperature. The resulting solution was stirred for 3 h at 25 °C. The crude product was purified by Prep-HPLC to afford the racemic product. The racemate was purified by chiral-HPLC to afford two enantiomers.

[(1R,2S)-2-cyanocyclopentyl] N-[8-amino-7-fluoro-6-(8-methyl-2,3-dihydro-1H-pyrido[2,3-b][1,4]oxazin-7-yl)-3-isoquinolyl]carbamate (C5) (28.4 mg,0.0614 mmol, 18.5% yield) as a yellow solid. Retention time: 1.726 min (CHIRALPAK IG, 0.46*5cm; 3 mm; Mobile Phase A: MtBE (0.1%DEA)-HPLC, Mobile Phase B: MeOH-HPLC; Flow rate: 1.0 mL/min; Gradient: 30 B to 30 B in 4 min; 254/220 nm). LCMS (ESI) $[M+H]^+$ = 463; $^1$H NMR (400 MHz, DMSO-$d_6$) δ 10.22 (s, 1H), 9.34 (s, 1H), 7.98 (s, 1H), 7.33 (s, 1H), 6.87 (d, $J$ = 6.1 Hz, 1H), 6.23 (s, 2H), 5.68 (s, 1H), 5.26–5.18 (m, 1H), 4.29 (t, $J$ = 4.4 Hz, 2H), 3.35 (s, 2H) 3.17 (dd, $J$ = 7.6, 4.9 Hz, 1H), 2.13 (ddt, $J$ = 24.2, 10.8, 5.5 Hz, 2H), 1.92 (d, $J$ = 1.7 Hz, 3H), 1.84-1.76 (dt, $J$ = 7.9, 5.2 Hz, 4H).

[(1S,2R)-2-cyanocyclopentyl] N-[8-amino-7-fluoro-6-(8-methyl-2,3-dihydro-1H-pyrido[2,3-b][1,4]oxazin-7-yl)-3-isoquinolyl]carbamate (enantiomer of C5) (26.6 mg, 0.0575 mmol, 17.3% yield) as a yellow solid. Retention time: 2.425 min. (CHIRALPAK IG, 0.46*5cm; 3 mm; Mobile Phase A: MtBE (0.1%DEA)-HPLC, Mobile Phase B: MeOH-HPLC; Flow rate: 1.0 mL/min; Gradient: 30 B to 30 B in 4 min; 254/220 nm). LCMS (ESI) $[M+H]^+$ = 463; $^1$H NMR (400 MHz, DMSO-$d_6$) δ 10.22 (s, 1H), 9.34 (s, 1H), 7.98 (s, 1H), 7.33 (s, 1H), 6.87 (d, $J$ = 6.1 Hz, 1H), 6.23 (s, 2H), 5.68 (s, 1H), 5.26–5.18 (m, 1H), 4.29 (t, $J$ = 4.4 Hz, 2H), 3.35 (s, 2H) 3.17 (dd, $J$ = 7.6, 4.9 Hz, 1H), 2.13 (ddt, $J$ = 24.2, 10.8, 5.5 Hz, 2H), 1.92 (d, $J$ = 1.7 Hz, 3H), 1.84-1.76 (dt, $J$ = 7.9, 5.2 Hz, 4H).

### 1.19.6 C6

Chemical name: 1-(3,5-difluoro-4-((3-(trifluoromethyl)-1H-pyrrolo[2,3-b]pyridin-4-yl)oxy)phenyl)-3-(3-morpholinopropyl)urea
Exact synthetic protocol for this compound is listed as "Example 1" in the patent: WO2018228920 A1
https://pubchem.ncbi.nlm.nih.gov/compound/137362730

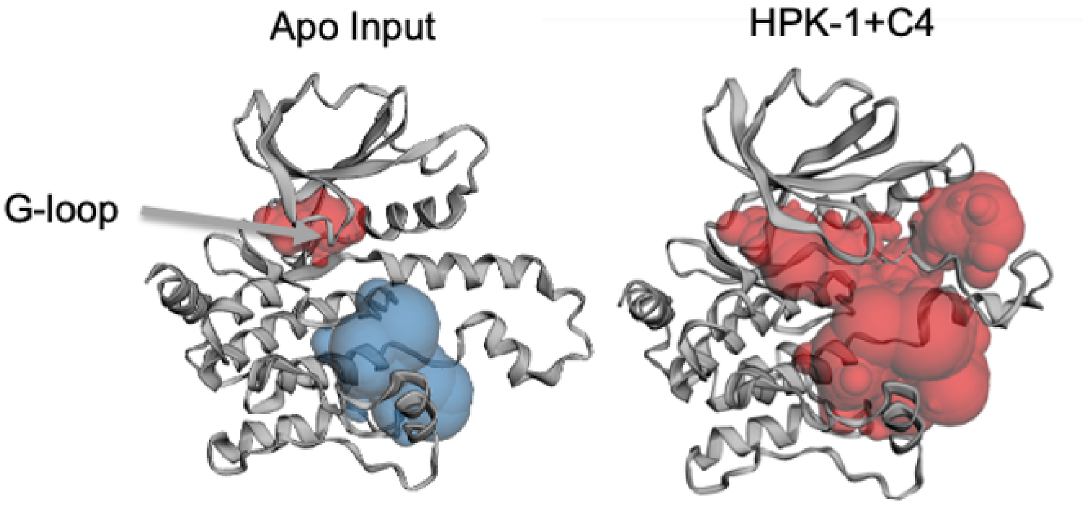


Figure 4: Apo HPK1 (left) starting structure has a collapsed binding pocket (red) with volume of approximately 143.9 Å3 and blue pocket with 859.2 Å$^3$ as assessed by CastP39 using a 1.4 Å probe radius. On the right, the same analysis was applied to HPK1 bound to C6 with the ligand removed. Due to extensive conformational changes involving backbone and sidechains, the larger pocket blue pocket in the apo structure merges with the binding pocket to create one large pocket (red, right) with a volume of 2737.6 Å3 (+272% as compared to the total volume of the two pockets in the Apo input).

### 2 Remapping Algorithm

#### 2.1 Aligning and mapping molecular structures

Comparing equivalent molecular structures of different origins is traditionally challenging due to the inconsistent assignment of atom names. The naming conventions for atoms from a crystal structure will be different from those generated from Gaussian, MOE, and other Pharmadesign-based software. This even extends to identically named molecules containing indistinguishable atoms on symmetric substructures (e.g., all carbon rings, methyl groups, etc.). As a result, much manual work is required to verify that the atomic contacts and their distances are equivalent across structures. Here, we have developed a method that maps the names of one atomic structure onto another and identifies degeneracies within the structure utilizing a graph-based substructure alignment algorithm. The context and solution of the issue are described in the following. Two chemically identical molecules generated from different software packages (and sometimes the same) will contain the same connections and atomic elements, but the names for the individual atoms (C1,C13,C20, etc.) will differ. The manual solution would be to overlay each structure in some visualization software and rename the atoms within one of the molecules to match the other. This method is effective, but becomes time-intensive when dozens of molecules are involved or when the molecules are large. To reduce the need for manual intervention, we developed an algorithm that correctly maps the atom names of one molecule onto another. The algorithm aligns two molecules' undirected atomic graph representations by determining equivalent ordered trees and the transformation between them. The algorithm is as follows:

1. construct a graph of each molecule comprised of atoms and their bonds.
2. construct weighted trees originating from each vertex on the graph
3. identify unique and degenerate atoms on each molecule
4. find equivalent trees between molecules

5. generate transformation matrix between molecules

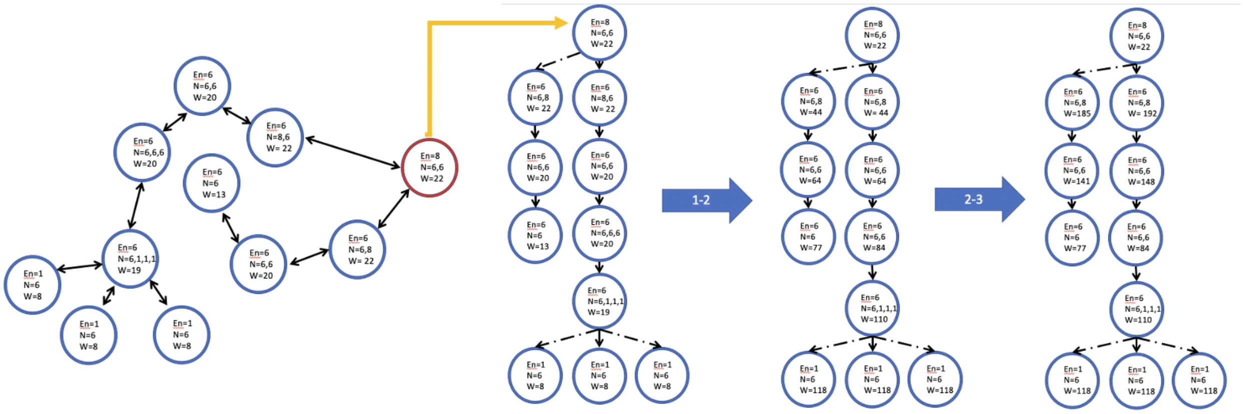


Figure 5: Ordered-Tree Construction

Graphs of the molecule are constructed by setting each atom as a vertex, then each of its bonds as a connection to another vertex. Each vertex contains additional information such as the atomic element number of the atom, the atomic element numbers of its neighbors, and a weight.

To construct a unique weighted tree from any arbitrary vertex on the graph, we utilize the weight identifier. The weight is calculated in three passes. In the first pass, the weight is assigned by the following sum: atomic elemental number of the target vertex + atomic elemental number of neighboring vertices + the number of children. Then we propagate the weights through to their descendants.

If at some depth, a tree contains vertices with the same weights, they are possibly degenerate. In that instance, we add the weights of their relatives until the weights are unique or until the tree terminates. This process is done in two passes: first, we step down the tree from the vertices of interest, and then up the tree if the weights are still degenerate. This does not guarantee that all vertices have a unique weight nor that unique vertices within a single a tree are truly unique; rather, it guarantees that all non-unique vertices are degenerate within the graph. These degeneracies are a result of symmetry between the vertices, making them identical and interchangeable (e.g., hydrogen atoms on a methyl group, atoms on all carbon rings, etc.).

From here onward, determining the transformation is trivial due to each globally unique vertex containing a corresponding weight to itself in the other graph. In the cases they are not unique, we are able to determine the groups of vertices that are symmetric and equivalent. A more rigorous implementation of this process can be seen below.

#### 2.2 Implementation

As a precursor, we introduce a non-standard notation of function inputs. $F(X, Y, Z; A, b, c)$ where all inputs preceding the ';' are static structures globally defined, and the superseding inputs are dynamic and definable at function call. In the cases $F(...; A, b, c)$ we compact the static portion of the inputs to save on white space. Vertex and edge are used in preference over atom and bond, respectively.

##### 2.2.1 Tree construction

Let $S$ be the ordered set of $N$ vertices

$$S = \{0, 1, 2, 3, ..., N - 1\}$$

Let $E$ be the list of elemental numbers corresponding to the vertices in $S$, such that $|S| = |E| = N$.

Let $\Xi$ by the ordered set of unique strings corresponding to vertex label of S

$$\Xi = \{``H1"\,``C1", ``C5", ``C3", ..., \xi_{n-1}\}$$

Let $\mathbf{C}$ be a connectivity matrix describing the network of edges in the vertices in $S$, where the column and row indices correspond to a unique element of S. It is important to note that $\mathbf{C}$ must be fully traversable, meaning one can construct a path between any two elements in the matrix.

Let us define a function, $F(\mathbf{C}; i) = F(i)$, that takes input $i \in S$ and returns an n-variable tuple comprised of connected elements to vertex value $i$, as described in $\mathbf{C}_i$.

Now let us define a rooted tree $T$ from an arbitrary starting vertex, $T_0 \in S$. The rest of the tree takes the recursive definition:

$$T_n = \left\{ F(i) - \bigcup_{w=0}^{n-1} T_w \,\middle|\, (i \in T_{n-1}), < \right\}$$

Afterwards, we create a list of weights defined by:

$$W = (W_i)_{i=0}^{|S|-1}, \quad W_i = |F(i)| + E_i + Z(i)$$

where $Z(i)$ is a function defined for summing the elemental numbers for connected vertexes to $i$. Now we define a set $A$ with the same structure as T with atomic elements defined as:

$$A_{i,n} = W_{T_{i,n}} + \sum_{j}^{|K|} K_j, \quad K = (W_x | x \in (F(T_{i,n}) \cap T_{i-1}))$$

Then we update W by assigning the weights in A to W through T:

$$W_{T_{i,j}} = A_{i,j}$$

##### 2.2.2 Resolving degeneracies

Now let us define $J$ for determining degeneracies in $A$

$$J = (J_i)_{i=0}^{|T|-1}, \quad J_i = \left( (n) \middle| (n \neq j) \wedge (A_{i,n} = A_{i,j}) \right)$$

Now we sum the relatives of $J_i$ to the degeneracies in $A_i$ until they are resolved or until the sum terminates. Let us first define a function $\theta(T, C, P; i, k)$ which will return descendants or ascendants vertices of $P$ at depth $k$. This can be observed as rotating the root to $P$ and choosing to look for descendants on the LHS or RHS of the tree.

$$\theta(T, C; P, i, k) = \left\{ \begin{array}{ll} \theta(...; \{F(n) \cap T_{i+1} | n \in P\} - P, i+1, k), & \text{if } i < k \\ P, & \text{if } i = k \\ \theta(...; \{F(n) \cap T_{i-1} | n \in P\} - P, i-1, k), & \text{if } i > k \end{array} \right\}$$

With that we can define a function $\psi(T, C, A; i, a, d)$ that will sum the weights of the relatives through till level d.

$$\psi(T, C, A; i, a, d) = A_{i,a} + \sum_{\beta = i + \beta \frac{d-i}{|d-i|}}^{|d-i|} \sum_{j=1}^{|\Omega|} \Omega_j, \quad \Omega = \{W_\phi | \phi \in \theta(...; \{T_{i,a}\}, i, \beta)\}$$

As mentioned above, we sum the descendant weights of degeneracies until they are no longer degenerate or until termination of the sum. We define $\Psi(T, C, A; P, D = \emptyset, i, d)$ as a function that will return the depths of degeneracy set P that will result in resolution of degeneracy. We will also define the function $\beta(L,i,d)$ that returns the elements of L that have the same descendant weight at depth d.

$$\beta(L,i,d) = \left\{ a \,\middle|\, (a, c \in L_i) \wedge (a \neq c) \wedge (\psi(...;i,a,d) = \psi(...;i,c,d)) \right\}$$

$$\Psi^+(T,C,A;P,D,i,d) = \left\{ \begin{array}{ll} \Psi(...;\beta(P,i,d), D \cup (x \cup (d)|x \in (P - \beta(P,i,d))), i, d+1), & \text{if } \beta(P,i,d) \neq \emptyset \text{ and } d \neq |A| \\ D & \text{else} \end{array} \right\}$$

$$\Psi^-(T,C,A;P,D,i,d) = \left\{ \begin{array}{ll} \Psi(...;\beta(P,i,d), D \cup (x \cup (d)|x \in (P - \beta(P,i,d))), i, d-1), & \text{if } \beta(P,i,d) \neq \emptyset \text{ and } d \neq -1 \\ D & \text{else} \end{array} \right\}$$

Now we apply these functions in sequence or in alternation. For this case we will be applying $\Psi^+$ then $\Psi^-$ in sequence.

$$\begin{aligned} L &= (L_i)_{i=0}^{|J|-1}, \quad L_i = \Psi^+(...;J_i,\emptyset,i) \\ U &= (U_i)_{i=0}^{|L|-1}, \quad U_i = J_i - \bigcup_{w=0}^{|L_i|-1} \{L_{i,w,0}\} \\ H &= (H_i)_{i=0}^{|U|-1}, \quad H_i = \Psi^-(...;U_i,\emptyset,i) \\ G &= (G_i)_{i=0}^{|H|-1}, \quad G_i = L_i \cup H_i \\ U' &= (U'_i)_{i=0}^{|L|-1}, \quad U'_i = J_i - \bigcup_{w=0}^{|G_i|-1} \{G_{i,w,0}\} \end{aligned}$$

The list $U'$ describes unsolvable degeneracies in the tree with origin at $T_0$. There lies an exception for this set to be incomplete, where $T_0$ is a degenerate vertex. However, we can resolve that issue by creating a set of trees containing every possible $T_0$. By comparing all the trees, we can determine all the degeneracies in all odd numbered systems. For an atom to be unique, it needs to be resolved in all trees, the inverse is also true for degenerate vertexes. For an atom to be degenerate, there must exist a tree where it is unresolved. A good example of this is the Water Molecule. if $T_0$ starts on either of the hydrogen atoms, the tree would be fully resolved. However, by starting on the Oxygen we would recognize the H's are degenerate.

In even numbered systems, like O2 or Dimethyl sulfoxide, the current method breaks down for identifying those symmetries. To resolve this aspect, we check for if the molecule contains even or odd number atoms. For even numbered systems, we add a fake atom vertex that has a connection with two atoms bonded to each other and use that fake atom at $T_0$ for solving the tree using the standard methodology above. By doing so we've converted an even numbered system into an odd one, which is solvable with the envisioned method.

##### 2.2.3 Mapping labels between structures

With the techniques above, we have all the information required for mapping labels between structures. Given two identical graphs with unique labels, we can relate them by their set of weighted trees. By taking the tree with the most symmetry from both structures we can identify that any vertex with the same weight at the same depth are equivalent. In the case of degeneracies, we make note of them, choose the mapping at random, and propagate that choice into any immediate relatives that are also degenerate.
